# Crystal structure and biochemical analysis suggest that YjoB ATPase is a substrate-specific molecular chaperone

**DOI:** 10.1101/2022.05.06.490860

**Authors:** Eunju Kwon, Pawan Dahal, Dong Young Kim

**Author notes:** These authors equally contributed to this work (Eunju Kwon, Pawan Dahal).

## Abstract

AAA+ ATPases are ubiquitous proteins associated with most cellular processes, including DNA unwinding and protein unfolding. Their functional and structural properties are typically determined by domains and motifs added to the conserved ATPases domain. Currently, the molecular function and structure of many ATPases remain elusive. Here, we report the crystal structure and biochemical analyses of YjoB, a *Bacillus subtilis* AAA+ protein. The crystal structure revealed that the YjoB hexamer forms a bucket hat-shaped structure with a porous chamber. Biochemical analyses showed that YjoB prevents the aggregation of vegetative catalase KatA and gluconeogenesis-specific glyceraldehyde-3 phosphate dehydrogenase GapB, but not citrate synthase, a conventional substrate. Structural and biochemical analyses further showed that the internal chamber of YjoB is necessary for the chaperone activity. Our results suggest that YjoB, conserved in the class *Bacilli*, is a molecular chaperone acting in the starvation/stationary phases of *B. subtilis* growth.

## INTRODUCTION

P-loop NTPases hydrolyze the β–γ phosphate bond of a bound nucleoside triphosphate, thereby causing conformational changes in their structure or in that of other molecules (1). As P-loop NTPases, AAA+ (ATPases associated with various cellular activities) superfamily proteins contain at least one ATPase (AAA) domain that hydrolyzes ATP and converts this biochemical energy into mechanical power (2–4). The AAA domains share conserved overall folds and characteristic motifs across all species (5, 6). Walker-A (GX_4_GK[S/T], where X is any amino acid) and Walker-B (hhhhD[D/E], where h is a hydrophobic residue) motifs are characteristics of typical P-loop NTPases, including AAA+ proteins. Nucleotide-interacting motifs that distinguish AAA+ proteins from other P-loop NTPases are sensor-1 (Asn required for the proper orientation of a water molecule during ATP hydrolysis), sensor-2 (Arg that interacts with the γ-phosphate of ATP), and arginine finger (Arg that transfers hydrolysis energy to neighboring subunits). The AAA domains are usually assembled into a ring-shaped hexamer or heptamer, which forms a central pore through which substrates can pass (4, 5).

In contrast to conserved hydrolytic functions, AAA+ proteins recognize specific substrates and perform diverse non-conserved cellular functions (7, 8). Many AAA+ proteins selectively unfold target proteins, which are subsequently subjected to degradation by a chambered protease (9), refolding by a co-chaperone (10, 11), or disassembly of a protein complex (12, 13). Some AAA+ proteins, such as DNA helicase, are involved in DNA segregation (8). The substrate selectivity of AAA+ proteins is usually influenced by additional motifs or domains that recognize substrates directly or through adaptor proteins. For example, ClpX ATPase recognizes some substrates through its N-terminal domain (NTD), which binds the substrate adaptor SspB (14–16). The diversity of substrates and the diverse cellular functions of many AAA+ proteins have not been fully characterized.

*Bacillus subtilis* YjoB consists of a single AAA domain and a unique NTD (17); the AAA domain is conserved similar to that in canonical AAA+ proteins, whereas its NTD exhibits no sequence similarity to other known proteins. As a member of the σ^W^ (sigma factor W) regulon, YjoB is overexpressed in the stationary phase (18) or in response to envelope stresses induced by alkaline pH, salt, and antibiotics (19–23). This suggests that YjoB may contribute to counteracting the stresses that activate σ^W^. However, the molecular function of YjoB has not been characterized. Here, we determined the crystal structure of the YjoB hexamer and elucidated its possible molecular role based on the identified YjoB-binding proteins and its specific activity.

## RESULTS

### Overall structure of the YjoB monomer

*B. subtilis* YjoB (residues 1-423) was overexpressed in *Escherichia coli* and purified using immobilized-metal affinity chromatography (IMAC) and size-exclusion chromatography (SEC). The purified YjoB exhibited ATPase activity with K_m_ of 13.6 ± 1.5 μM and V_max_ of 228.7 ± 6.3 nmol/min/mg, whereas its Walker-B mutants, E280Q and D281N, showed no activity (Fig. 1A). For structure determination, an initial electron density map was calculated using the selenomethionine (SeMet)-labeled YjoB via the multiple-wavelength anomalous dispersion (MAD) phasing method. Two chains were found in the asymmetric unit (Supplementary Fig. S1A), and each chain was traced into an electron density map in the residue ranges 4–322/327–404 and 3– 404, covering the whole core structure. The crystal structure of the native YjoB was determined at 2.62 Å resolution with a R/R_free_ value of 20.0/23.6% (Supplementary Fig. S1B and Supplementary Table S1).

**Figure 1.**
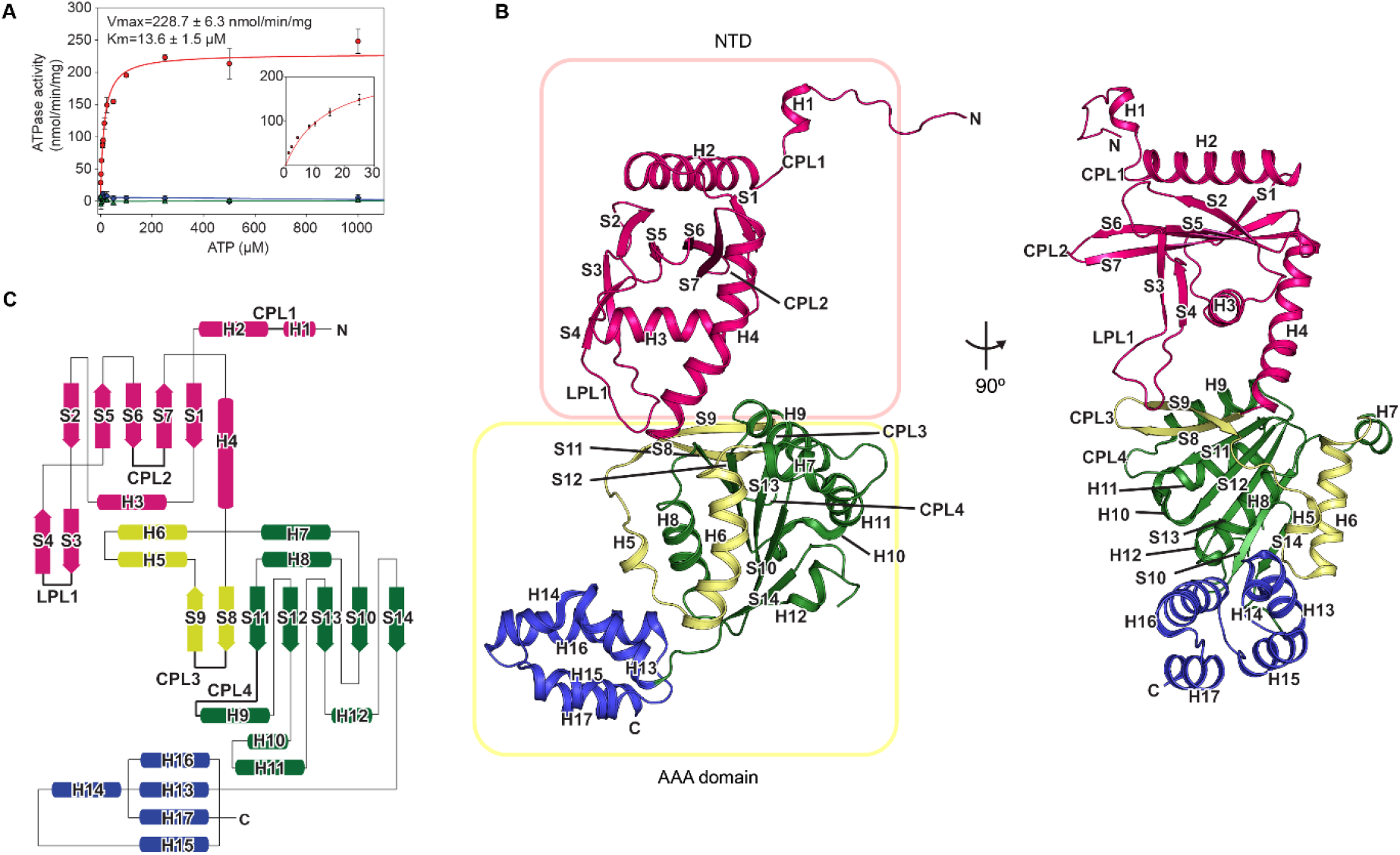
Overall structure of the YjoB monomer. (**A**) ATPase activity of YjoB. (**B**) Ribbon diagram and (**C**) topology model of the YjoB monomer. Secondary structures are denoted as H1– H17 for helices and S1–S14 for strands. CPL1–CPL4 and LPL1 represent the central and lateral pore loops, respectively. NTD is displayed in magenta. Core α/β fold, insertion motifs (S8–S9 and H5–H6), and C-terminal helical bundle of the AAA domain are shown in green, light green, and blue, respectively.

The YjoB monomer in the crystal structure consists of an NTD (residues 1–159) and an AAA domain (residues 160–404) (Fig. 1B, 1C and Supplementary Fig. S2). The YjoB-AAA domain forms a core α/β fold consisting of a parallel β-sheet (strand order of S11, S12, S13, S10, and S14) and peripheral helices (H7–H12), which is a characteristic fold of AAA+ proteins (Fig. 1B, 1C). The five C-terminal helices (H13–H17) form a helical bundle independent of the core α/β fold. A two-stranded antiparallel β-sheet (S8 and S9) and two helices (H5 and H6) are placed at the N-terminus of the AAA domain. In structural homology searches, the overall fold of the YjoB-AAA domain was similar to that of the classical AAA clade of AAA+ proteins (6). In particular, the AAA domains of the proteasomal subunit Rpt4, the protein unfoldase CDC48, and the mitochondrial chaperone Bcs1, which belong to the classical AAA clade, were superimposed onto the YjoB-AAA domain, with low root-mean-square deviation (RMSD) values of 2.8 Å for 229 Cα atoms, 2.4 Å for 220 Cα atoms, and 3.0 Å for 210 Cα atoms (Supplementary Fig. S3A–S3C).

The YjoB-NTD forms a core structure with an antiparallel β-sheet (strand order of S1, S7, S6, S5, and S2) surrounded by a two-stranded β-sheet (S3 and S4) and three α-helices (H2–H4). In addition to the core structure, its N-terminus is extended by a short 3^10^-helix (H1) and an unstructured region (Fig. 1B, 1C). No YjoB-NTD-like proteins or domains were found in structural homology searches using the DALI server (24). However, the overall fold of the YjoB-NTD was similar to that of the Bcs1-NTD, although they do not share a significant sequence identity. Only four of the 159 residues in YjoB-NTD aligned with *Saccharomyces cerevisiae* Bcs1-NTD as identical residues in a structure-based sequence alignment (Supplementary Fig. S3D). The core structure of YjoB-NTD was superimposed on the Bcs1-NTD structures from *S. cerevisiae* (PDB ID: 6SH3) and *Mus. musculus* (PDB ID: 6UKS) with RMSD values of 3.6 Å for 97 Cα atoms and 3.8 Å for 96 Cα atoms. A major structural difference between the NTD structures of YjoB and Bcs1 was found in the N-terminal helices (H1/H2 of YjoB and transmembrane helix H1 of Bcs1) (Supplementary Fig. S3E–S3I).

### Structure of the YjoB hexamer

The asymmetric unit of the YjoB crystal contains two YjoB monomers (Supplementary Fig. S1). Many residues distributed throughout the monomer participate in dimer interactions (Supplementary Table S2). The structure of the YjoB hexamer is formed by a 3-fold crystallographic symmetry of the asymmetric dimers, resulting in a large surface area burial (39,120 Å^2^ in a total of 98,650 Å^2^) and a significantly decreased ΔG (−143.6 kcal/mol) as calculated using the PISA program (25) (Fig. 2A). Consistently, YjoB was eluted in the volume between ferritin (440 kDa) and aldolase (158 kDa) in SEC (Supplementary Fig. S4A) and calculated as a hexamer by SEC with multi-angle laser light scattering (MALS) (Supplementary Fig. S4B). These observations show that YjoB assembles into a stable ring-shaped hexamer.

**Figure 2.**
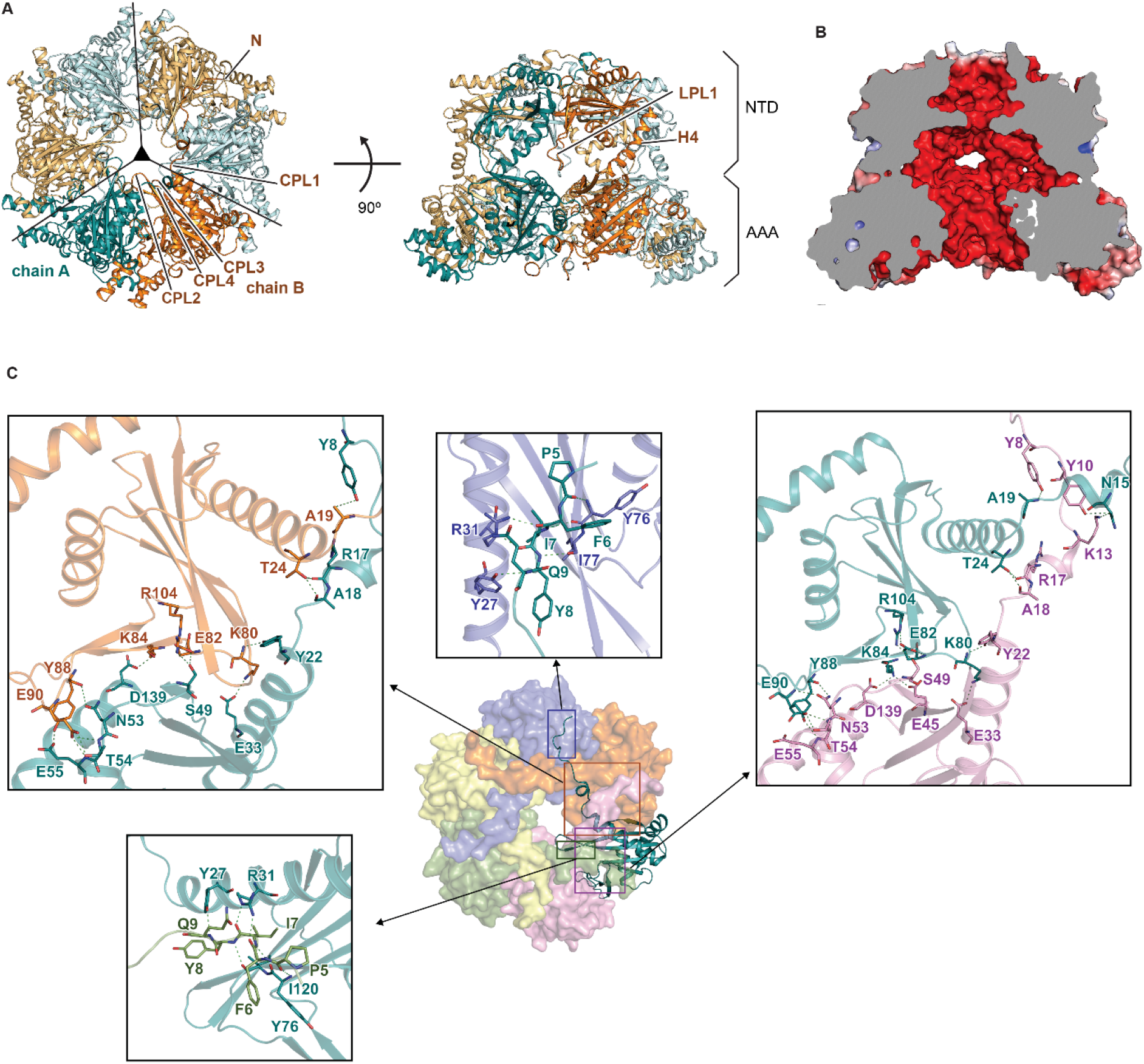
Crystal structure of the YjoB hexamer. (**A**) Top (left) and side (right) views of the YjoB hexamer. The two subunits in the asymmetric unit are shown in cyan and orange. The subunits generated by 3-fold crystallographic symmetry are shown in light cyan and light orange. (**B**) Charge distribution on the internal surfaces of the YjoB hexamer. Electrostatic potentials, from red (−5 kT/e) to blue (+5 kT/e), are plotted on solvent-accessible surfaces. (**C**) Binding interface for NTD hexamer assembly. One NTD and the other five NTD monomers in the NTD hexamer are represented by ribbon and surface models, respectively. Subunits are colored differently. Binding interfaces between NTDs around the ribbon model are shown in the boxes.

In the crystal structure of the YjoB hexamer, the NTDs assemble into a compact and stable hexameric ring with a surface area burial of 24,120 Å^2^ in a total surface of 42,870 Å^2^ and ΔG of - 122.1 kcal/mol. The globular core of the NTD interacts with the neighboring NTD cores, and the extended N-terminus containing H1 and an unstructured region interacts with the other three NTDs by protruding counterclockwise, forming an interlocking hexamer in which one NTD interacts with the other four NTDs (Fig. 2 and Supplementary Fig S4C, S4D). The YjoB-AAA domain also assembles into a hexamer, but it appears to be less stable than that of YjoB-NTD (Fig. S4E, S4F). The surface area burial and ΔG by the hexameric assembly of the YjoB-AAA domain are calculated to be 13,570 Å^2^ in a total surface of 63,659 Å^2^ and −24.2 kcal/mol, respectively. Moreover, the YjoB-AAA domain was eluted as an oligomer in SEC but was slightly heterogeneous depending on buffer conditions (Supplementary Fig. S4E). Thus, the crystal structure of YjoB shows that both NTD and AAA domains independently assemble into hexameric rings.

In the YjoB hexamer structure, a short 3^10^-helix H1, CPL1 (a central pore loop between H1 and H2; 18-AAAG-21), and CPL2 (between S6 and S7; 124-QDG-126) are aligned along the central pore of the NTD hexamer. CPL3 (between S8 and S9; 167-GDGG-170) and CPL4 (between S11 and H9; 256-FTS-258) are aligned along the central pore of the AAA hexamer (Figs. 1 and 2). The pore sizes of the NTD and AAA domain in the YjoB hexamer are restricted to 10 and 16 Å in diameter, respectively, by negatively charged residues, D125 in CPL2 and D168 in CPL3 (Supplementary Fig. S5A).

In the YjoB-NTD structure, the H4 helix and LPL1 (a lateral pore loop between S3 and S4; residues 88-YDEDEKEPDY-97) are placed outside of the NTD core, similar to jellyfish tentacles (Fig. 2A). As the hemispherical NTD hexamer is placed on the AAA hexamer, the YjoB hexamer exhibits a bucket hat-shaped structure harboring a porous chamber with approximate dimensions of 40 × 40 × 25 (Å) (Fig. 2B and Supplementary Figs. S5 and S6). The chamber volume is calculated to be approximately 2.6 × 10^5^ Å^3^, corresponding to a protein size of 21 kDa (26, 27). In addition to the central pore formed by a hexameric assembly of YjoB, the internal chamber has lateral pores formed by H3, H4, and LPL1 of the NTDs (Supplementary Fig. S5). The chamber surface, containing the area of central and lateral pores, is highly negatively charged, whereas the outer surface of the YjoB hexamer is relatively less hydrophilic (Fig. 2B and Supplementary Fig. S6A).

### Structure of the ADP-binding form of YjoB

To determine the crystal structure of the nucleotide-binding forms, YjoB was mixed with a 10-fold excess of ADP, AMPPNP, or ATPγS and was then crystallized. The electron density map of ADP was observed in only one of two YjoB chains in the crystal structure of YjoB mixed with ADP, whereas no additional density map of an adenine nucleotide was observed in the crystal structures of YjoB mixed with nonhydrolyzable ATP analogs (AMPPNP and ATPγS). Therefore, only the crystal structure of ADP-bound YjoB (YjoB-ADP) was determined at 2.70 Å resolution (Supplementary Table S1).

The structure of the canonical AAA+ protein shows that ADP binds to Walker-A and sensor-2 motifs (4). In the crystal structure of YjoB-ADP, the phosphates of ADP interact with residues 230–235 of Walker-A through hydrogen bonds, and the base of ADP interacts with F373 in sensor-2 through hydrophobic interactions. Walker-B, sensor-1, and Arg-finger do not participate in the ADP interaction, as seen in the structure of other AAA+ proteins that bind ADP (28, 29) (Fig. 3 and Supplementary Fig. S2). The structure of YjoB-ADP in the asymmetric unit was superimposed onto those of native YjoB with RMSD values of 0.4 Å for 797 Cα atoms, indicating that no significant conformational changes were observed between the structures of native YjoB and YjoB-ADP. Two monomers in the asymmetric unit of YjoB-ADP (apo and ADP-binding forms) were superimposed on each other with RMSD values of 1.9 Å for 396 Cα atoms. This shows that the conformational difference between the two monomers in an asymmetric unit is larger than the conformational change induced by ADP binding (Fig. 3D and Supplementary Fig. S7).

**Figure 3.**
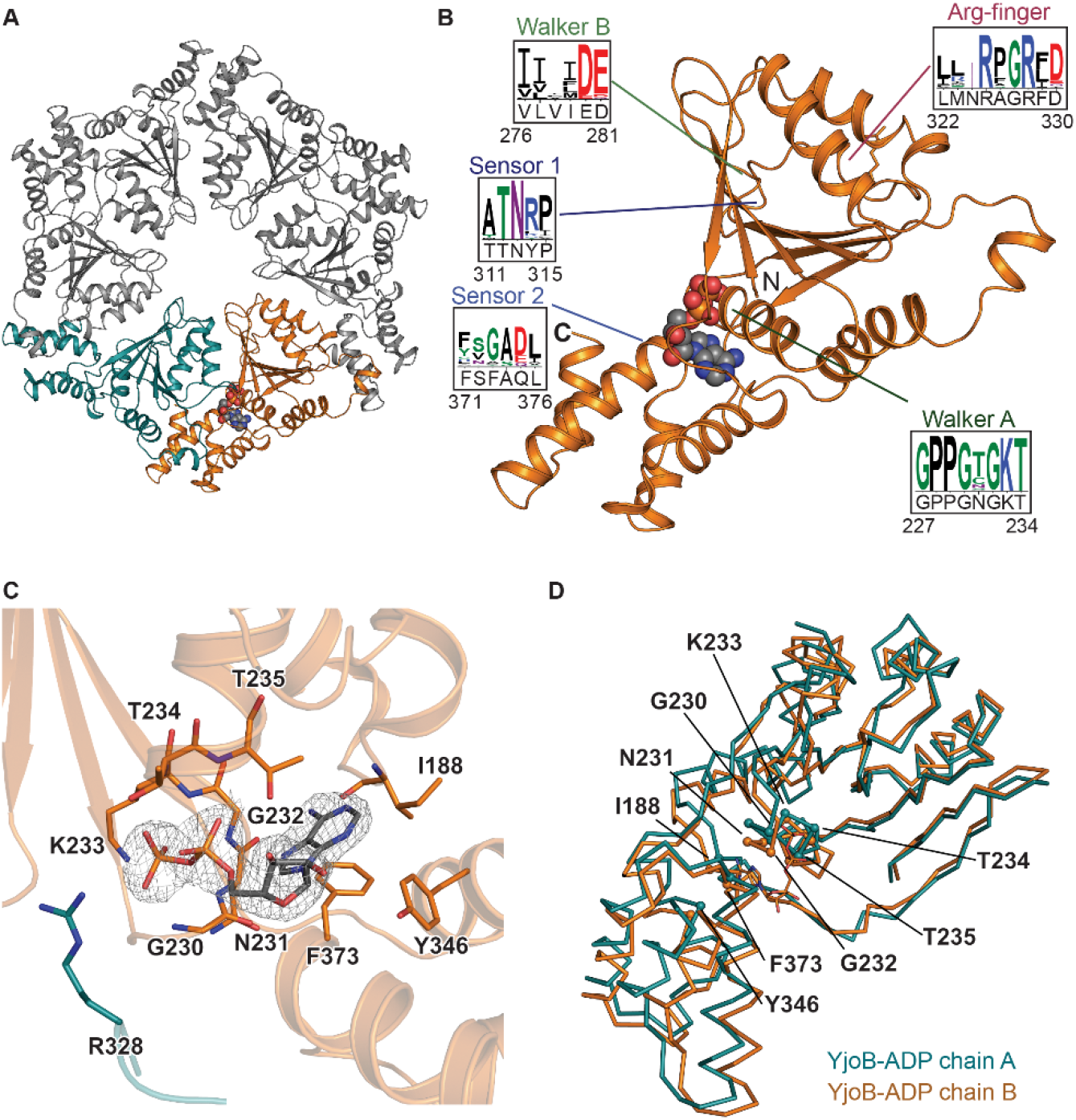
Structure of AAA domain in the crystal structure of YjoB-ADP. (**A**) Ribbon diagram of the YjoB-AAA hexamer. The two AAA domains in the asymmetric unit are shown in cyan and orange. ADP is represented by a sphere model. (**B**) Structure of the YjoB-AAA domain that binds ADP. Motifs conserved among AAA+ proteins are represented with the sequence logos. The sequence logo was prepared based on the sequence alignment of AAA+ proteins whose structures have been deposited in the PDB. (**C**) Interactions between AAA and ADP. The electron density of ADP is shown as a gray mesh at a contour level of 1.5. Residues around ADP are shown as stick models. R328 in the arginine finger of the neighboring subunit is shown as a red stick model. (**D**) Structural comparison between two AAAs in the asymmetric unit of YjoB-ADP crystal. The structures of the two AAA domains are superimposed and represented by Cα trace models. The AAA structures in the apo- and ADP-binding forms are shown in cyan and orange, respectively.

### YjoB suppresses aggregation of KatA and GapB

As the function of YjoB is unknown, we attempted to identify YjoB-binding proteins using affinity purification-mass spectroscopy (AP-MS) as a first approach to determine its function. Recombinant YjoB with N-terminal FLAG and Strep affinity tags (FLAG-Strep-YjoB) was expressed in *B. subtilis* and purified using Strep-pulldown and subsequent FLAG-immunoprecipitation. Proteins eluted with YjoB were then analyzed using liquid chromatography-tandem mass spectroscopy (LC-MS/MS). A total of 103 YjoB-binding proteins were identified in the volcano plot using EGFP-binding proteins as a control, of which eleven proteins had values higher than *p* = 0.05 and Log_2_(fold-change) = 5.0 (Fig. 4A). Among the eleven proteins, GapB, KatA, SufA, TrxA, YjoA, YphP, and YutI showed higher values than other proteins in emPAI (exponentially modified protein abundance index level) indicating protein abundance (30) (Fig. 4B). GapB is an NADP^+^-dependent glyceraldehyde-3 phosphate dehydrogenase. KatA is a catalase that binds heme as a cofactor (31, 32). SufA and YutI are putative scaffold proteins that bind iron-sulfur clusters (Fe–S) for the assembly of Fe-S and target proteins. TrxA is a thioredoxin. YjoA is a DinB-fold protein with a metal-binding triad with an unknown function. YphP is a putative disulfide isomerase.

**Figure 4.**
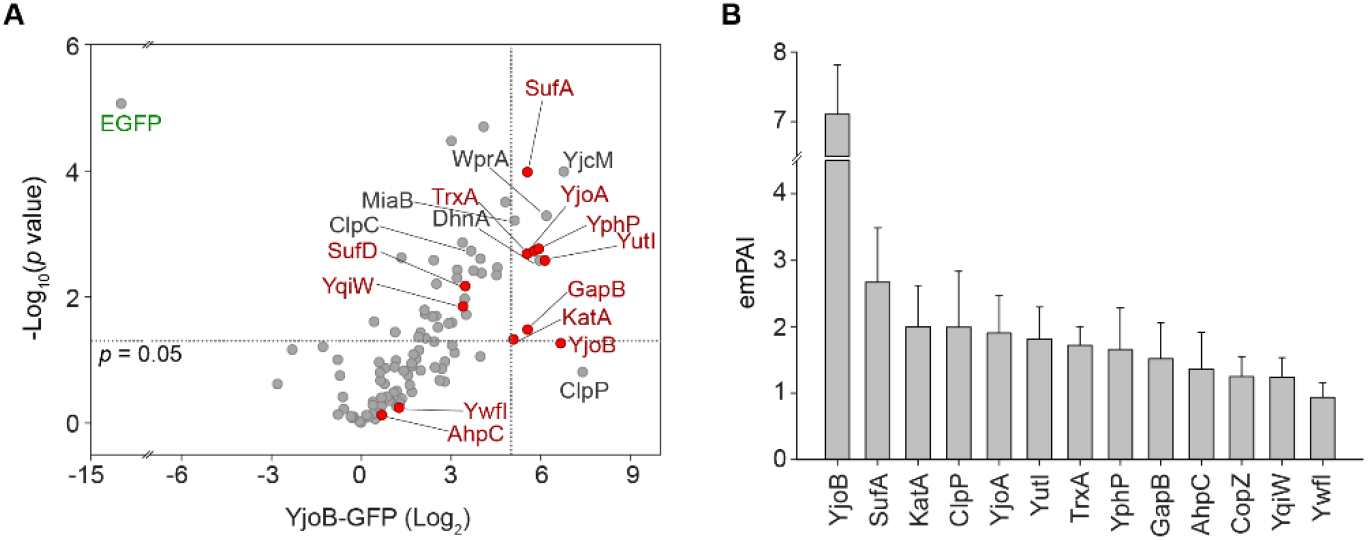
LC-MS/MS analysis of YjoB-binding proteins. (**A**) Volcano plot of 105 proteins identified as YjoB- and EGFP-biding proteins. A total of 103 YjoB-binding proteins were identified. The significance (non-adjusted *p*-value; -Log_10_(*p*-value)) and fold-change (Log_2_(fold-change)) of each protein are plotted on the x and y axes of the volcano plot. Horizontal and vertical dotted lines indicate *p*-value = 0.05 and fold-change = 5.0, respectively. Proteins marked in red were purified as substrate candidates for the measurement of YjoB chaperone activity. (**B**) emPAI values of YjoB-binding proteins. A total of 82 proteins were identified in three independent experiments. The top 13 values are presented as the mean ± standard error of three replicates.

To assess the chaperone activity of YjoB, YjoB-binding proteins were purified, and their heat-induced aggregation was examined to find suitable substrates for measuring chaperone activity (Supplementary Fig. S8A, S8B). A total of 11 YjoB-binding proteins were purified: GapB, KatA, SufA, TrxA, YjoA, YphP, YutI, and the other binding proteins (AhpC, SufD, YqiW, and YwfI). Of these, KatA (54.8 kDa) and GapB (37.5 KDa) formed aggregates at 42 °C, whereas the other proteins were heat-stable even at 64 °C (Supplementary Fig. S8A). During protein purification, KatA was eluted in three different oligomeric forms on SEC (Supplementary Fig. S8C). KatA was mostly in a monomeric form (KatA-1), whereas a small fraction was estimated to be a dimer (KatA-2) and higher oligomers (KatA-3) (Supplementary Fig. S8C, S8D). KatA-2 and KatA-3 showed absorbance peaks at 410 nm, indicating that they contain heme as a cofactor. KatA-1 did not show significant absorbance (Supplementary Fig. S8E) and was easily heat-aggregated at 42 °C (Supplementary Fig. S8A, S8B).

In the measurement of chaperone activity, aggregation of KatA-1 and GapB was suppressed in a dose-dependent manner by YjoB at 42 °C and was not prevented by bovine serum albumin (BSA) (Fig. 5A, 5B, and Supplementary Fig. S9C–S9F). Aggregation of KatA-2 and KatA-3 was also suppressed by YjoB at 64 °C, but the effect was limited owing to the relative thermal stability of the KatA oligomers (Supplementary Fig. S9A, S9B). In contrast to the YjoB-mediated inhibition of KatA-1 and GapB aggregation, YjoB did not prevent the aggregation of citrate synthase (CS), a conventional substrate for measuring chaperone activity (Fig. 5C). This differs from GroEL chaperonin preventing all aggregation of KatA-1, GapB, and CS (Fig. 5A–5C, and Supplementary Fig. S9). Therefore, YjoB appears to suppress aggregation in a substrate-specific manner.

**Figure 5.**
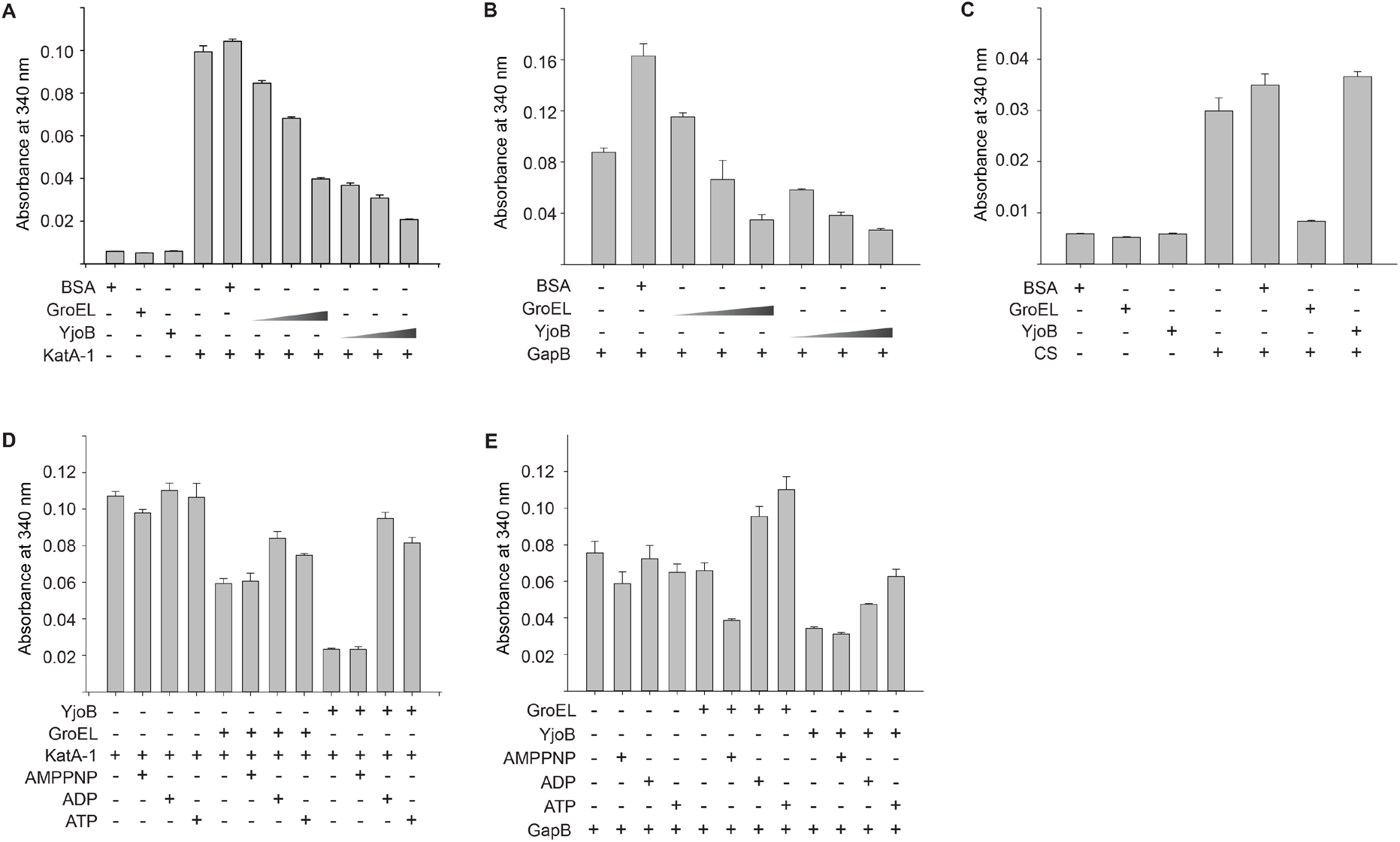
Chaperone activity of YjoB. (**A–C**) Thermal aggregation of KatA-1 (**A**), GapB (**B**), and CS (**C**) in the presence of YjoB or GroEL. (**D, E**) Thermal aggregation of KatA-1 (**D**) and GapB (**E**) in the presence of YjoB and adenine nucleotides. Bar diagrams in (**A–E**) show the level of aggregation at the 150 min time point in the absorbance spectra in Supplementary Figs. S9C–S9H and S10A**–**S10F. Absorbance was measured at 42 °C. All data in (**A–E**) are presented as the mean ± standard error of three replicates.

To investigate whether the suppression effect of YjoB is nucleotide-dependent, the activity was measured in the presence of ATP, ADP, or AMPPNP. Compared with the activity in the absence of the nucleotide, AMPPNP either did not change or slightly reduced aggregation of KatA-1 and GapB, whereas both ATP and ADP increased the aggregation (Fig. 5D, 5E and Supplementary Fig. S10A–S10F). The aggregation pattern by adenine nucleotides was similar to that of GroEL, which increased in the presence of ADP or ATP (Fig. 5D, 5E and Supplementary Fig. S10C, S10F–S10H). The Walker-B mutants of YjoB (E280Q and D281N), which do not hydrolyze ATP, also prevented KatA-1 aggregation at a level similar to that of YjoB in the absence of nucleotides (Supplementary Fig. S10I, S10J). Consistent with this, in SEC, KatA-1 aggregates were eluted separately from YjoB in the presence of ATP or ADP, whereas they coeluted with YjoB in the presence of AMPPNP or the absence of nucleotide (Supplementary Fig. S11A–S11C). These results suggest that YjoB binds a substrate in the ATP-binding state or the absence of nucleotides and releases it under ATP hydrolysis or ADP binding.

### Internal chamber is necessary for the chaperone activity of YjoB

The NTD and AAA domains of YjoB were purified to determine the domains responsible for chaperone activity. The YjoB-AAA domain was purified as a homogenous hexamer, but the oligomer was destabilized depending on buffer conditions (Supplementary Fig. S4E). The YjoB-NTD was separated into an oligomer (NTD-1) and a monomer (NTD-2) that are not in equilibrium (Supplementary Fig. S4F). In the activity assays, the NTD-2 and the AAA domain suppressed thermal aggregation of KatA at a lower degree than full-length YjoB, whereas NTD-1 did not, indicating that NTD-1 is close to an inactive form (Fig. 6A and Supplementary Fig. S12A).

**Figure 6.**
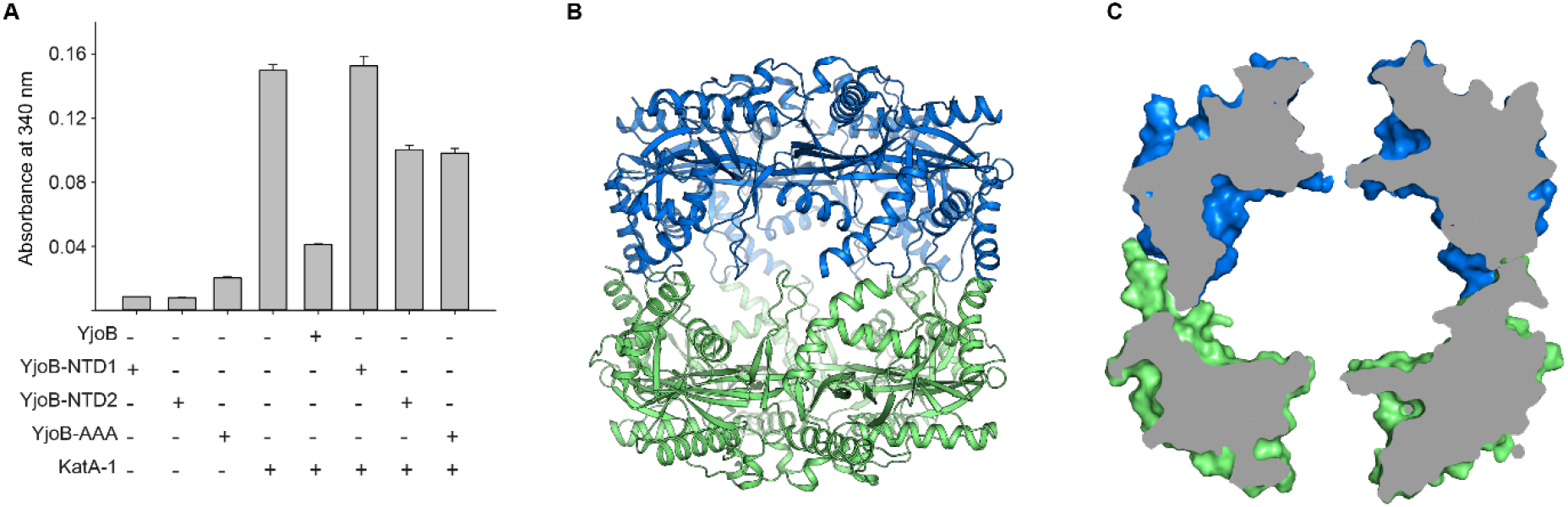
Role of the internal chamber in YjoB chaperone activity. (**A**) Thermal aggregation of KatA-1 in the presence of YjoB domains. YjoB-NTD was purified as an oligomer (NTD-1) and a monomer (NTD-2). Aggregation of KatA-1 (10 μM) was compared in the presence of YjoB, NTD-1, NTD-2, and YjoB-AAA (5 μM each). Bar diagrams show the level of KatA-1 aggregation at the 150 min time point in the absorbance spectra in Supplementary Fig. S12A. Absorbance was measured at 42 °C. Data from three replicates are presented as the mean ± standard error. (**B, C**) Crystal structure of the NTD-1 dodecamer. (**C**) The cross-sectional surface model shows that the internal chamber of NTD-1 is not accessible to the substrate. The two hexamer units of the NTD-1 are shown in different colors.

To examine why NTD-1 shows no chaperone activity, we crystallized NTD-1 and determined its crystal structure at 3.0 Å resolution (Supplementary Table S1). In the crystal structure, NTD-1 assembles into a stable ball-shaped dodecamer by dimerization of the NTD-1 hexamer, and the lateral pores are closed (Fig. 6B, 6C and Supplementary Fig. S12B). The structure shows that inactive NTD-1 restricts substrate access to the internal chamber, suggesting that exposure of the internal chamber to a substrate is required for chaperone activity of YjoB.

### YjoB enhances the catalase activity of KatA

KatA is a major vegetative catalase of *B. subtilis* that binds heme and decomposes hydrogen peroxide (H_2_O_2_) into water and oxygen (31, 32). As YjoB significantly prevented KatA aggregation, we next investigated whether YjoB affects the catalase activity of KatA.

In the catalase activity assays, inactive KatA-1 was activated by the addition of hemin at 30 °C (Fig. 7A and Supplementary Fig. S13A), indicating that hemin can be spontaneously incorporated into KatA-1. KatA-1 aggregation was also reduced in the presence of hemin, indicating that hemin is required for KatA stability (Supplementary Fig. S11E). The catalase activity of KatA-1 was further increased in the presence of ATP (Fig. 7A and Supplementary Fig. S13A). ATP itself appears to promote heme incorporation into KatA-1. In contrast to the activity enhancement by ATP, YjoB did not significantly affect KatA-1 activity at 30 °C, suggesting that YjoB does not affect KatA-1 activity under conditions that do not induce aggregation (Fig. 7A and Supplementary Fig. S13A). To examine whether YjoB enhances KatA-1 activity under KatA-1 misfolding conditions, KatA-1 was incubated with YjoB at 42 °C, and catalase activity was measured after incubation with hemin (Fig. 7A and Supplementary Fig. S13B). Compared with the activity measured at 30 °C, the catalase activity of KatA-1 after thermal incubation decreased in the presence of ATP and hemin, whereas it was maintained in the presence of YjoB (Fig. 7A and Supplementary Fig. S13A, S13B). This demonstrates that YjoB enhances KatA-1 activity by preventing its aggregation or misfolding. However, the ATP hydrolysis effect of YjoB could not be observed because ATP itself enhanced catalase activity. YjoB slightly enhanced the catalase activity of KatA-1 during the renaturation of denatured KatA-1 (Supplementary Fig. S13C, S13D).

**Figure 7.**
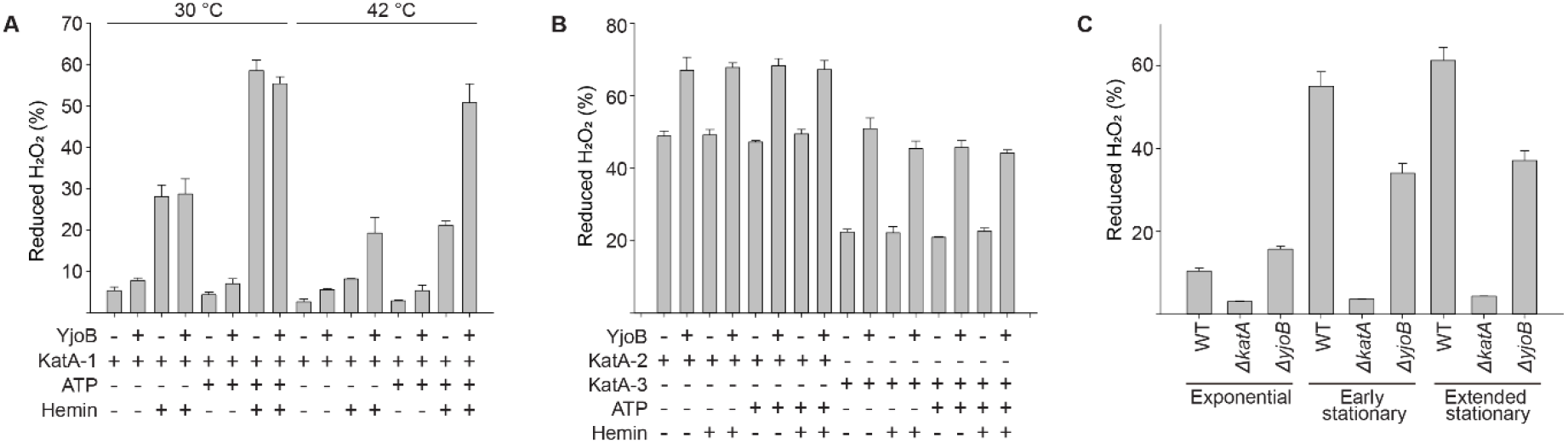
Catalase activity of KatA in the presence of YjoB. (**A**) Catalase activity of KatA-1. Mixtures of KatA-1 and YjoB were incubated at 30 °C or 42 °C, followed by further incubation with hemin at 30 °C. Decomposition of 20 mM H_2_O_2_ by 25 nM KatA-1 was measured at 30 °C. Bar diagrams show the values at the 20 min time point in the H_2_O_2_ decomposition curves in Supplementary Fig. S13A, S13B. (**B**) Catalase activity of KatA-2 and KatA-3. Mixtures of KatA-2 (or KatA-3) and YjoB were incubated at 42 °C, followed by further incubation with hemin at 30 °C. Decomposition of 20 mM H_2_O_2_ by 0.25 nM KatA-2 (or 0.25 nM KatA-3) was measured at 30 °C. Bar diagrams show the values at the 20 min time point in the H_2_O_2_ decomposition curves in Supplementary Fig. S13E, S13F. Data in (a–b) are presented as the mean ± standard error of three replicates. (**C**) Decomposition of 20 mM H_2_O_2_ by *B. subtilis* and its mutants (*ΔkatA* and *ΔyjoB*). Cells were harvested at the exponential, early stationary, and extended stationary growth phases. Bar diagrams show the values at the 20 min time point in the H_2_O_2_ decomposition curves in Supplementary Fig. S13G–S13I. Data are presented as the mean ± standard error of six replicates.

In contrast to KatA-1, the catalase activity of KatA-2 and KatA-3 was not enhanced by hemin and ATP, indicating that they are heme-containing active forms (Fig. 7B and Supplementary Figs. S8E and S13E, S13F). To examine whether YjoB enhances the activity of KatA-2 and KatA-3, their catalase activity was measured after incubation with YjoB at 42 °C. YjoB increased KatA-2 and KatA-3 activities by 1.4-fold and 2.5-fold, respectively, regardless of the presence of ATP or hemin (Fig. 7B and Supplementary Fig. S13E, S13F), although aggregation of KatA-2 and KatA-3 was not observed at 42 °C (Supplementary Fig. S8B). It is unclear why the catalase activity of KatA-2 and KatA-3 is enhanced by YjoB. YjoB may contribute to slight conformational changes in KatA-2 and KatA-3 that affects the catalase activity.

Expression of YjoB and KatA is induced in the stationary phase of *B. subtilis* growth (18, 31, 32). To examine whether YjoB enhances KatA activity in the stationary phase, we compared the catalase activity of wild-type *B. subtilis* and a *yjoB* deletion mutant collected at different bacterial growth phases. Catalase activity in wild-type *B. subtilis* remained low in the exponential phase (Fig. 7C and Supplementary Fig. S13G), but increased 6-fold during the early stationary phase and was maintained in the extended stationary phase (Fig. 7C and Supplementary Fig. S13H, S13I). Catalase activity of the *yjoB* deletion mutant doubled in the early stationary phase. Compared to wild-type *B. subtilis*, the *yjoB* deletion mutant exhibited approximately 30% higher catalase activity in the exponential phase but half the activity in the early stationary phase. (Fig. 7C and Supplementary Fig. S13G–S13I). These results suggest that YjoB enhances KatA activity during the stationary phase in which KatA is overexpressed.

## DISCUSSION

YjoB is an uncharacterized AAA+ ATPase first identified in the genome of *B. subtilis* (33). This study presents the crystal structure of YjoB and suggests its molecular function based on the relationship between YjoB and identified substrates. Our results show that *B. subtilis* YjoB may function as a substrate-specific molecular chaperone.

YjoB homologs with more than 50% sequence identity are found in the class *Bacilli*, mainly in the genera *Paenibacillus* and *Bacillus* of the order *Bacillales* (Supplementary Fig. S14). Other than the bacteria, plants (the genus *Arabidopsis*) and plant-infecting oomycetes (the genus *Phytophthora*) infrequently contain YjoB homologs as a domain of large proteins (Supplementary Fig. S2). Based on the sequence alignment of YjoB homologs, we classify YjoB as an AAA+ protein specifically conserved in the order *Bacillales*.

The overall fold of the YjoB-AAA domain is similar to that of proteasomal AAA subunits, the second AAA domain in Cdc48, and the AAA domain of mitochondrial Bcs1 (Bcs1-AAA domain). Notably, the YjoB-AAA domain is phylogenetically closest to the Bcs1-AAA domain among proteins with known structures (Supplementary Fig. S15). Both YjoB and Bcs1 have characteristic motifs conserved in classical AAA proteins but exceptionally, contain an ED sequence in the Walker-B (276-VLVI<u>ED</u>-281 in YjoB) motif instead of a DE sequence. A Glu residue in the Walker-B motif of classical AAA proteins primes a water molecule for a nucleophilic attack on the γ-phosphate of ATP (6). Although YjoB and Bcs1 contain Asp instead of Glu in the Walker-B motif, YjoB showed ATPase activity and Bcs1 changed its conformation in a nucleotide-dependent manner (34), indicating that they are active ATPases. This shows that the YjoB-AAA domain is an active ATPase similar to that in the Bcs1 family of the classical AAA clade.

Bcs1 function is involved in the integrity of the mitochondrial respiration complex III (35, 36). Rip1, a subunit of the cytochrome bc1 complex, is complexed with the Fe-S cluster in the matrix, translocated to the inner membrane via the internal chamber of Bcs1, and inserted into a cytochrome bc1 complex of the inner membrane (37). The overall shape of the Bcs1 heptamer is similar to that of the YjoB hexamer (Supplementary Figs. S5 and S6). The similarity in the overall shape of oligomeric forms between YjoB and Bcs1 suggests that they might partially share a reaction mechanism, such as using an internal chamber for their activity. A notable difference between YjoB and Bcs1 is that Bcs1 forms a heptameric ring-shaped structure and is anchored to the inner membrane via its N-terminal helix (34, 38), whereas YjoB assembles into a stable hexamer interlocked by its NTD and is a cytoplasmic protein with no transmembrane or amphiphilic helices (Supplementary Figs. S4C, S4D and S6). Another difference between YjoB and Bcs1 is the charge distribution on their internal chamber. The internal chamber of YjoB is highly negatively charged, whereas that of Bcs1 does not exhibit a high degree of charge distribution (Fig. 2B and Supplementary Fig. S6C).

In our studies, YjoB suppressed the aggregation of KatA and GapB. In *B. subtilis*, YjoB expression is upregulated by SigW which is activated when bacterial growth has reached the stationary phase or in response to envelope stress (18–23). KatA expression is induced by glucose starvation, in the stationary phase, and during oxidative stress (31, 32). GapB expression is induced by glucose starvation conditions that promote gluconeogenesis (39). These suggest that YjoB may be a chaperone that enhances the integrity of specific proteins during the stationary/starvation phase. Consistent with this, the catalase activity increased during the stationary phase of wild-type *B. subtilis*, whereas that in *yjoB* deletion mutant was restricted (Fig. 7C and Supplementary Fig. S13G–S13I).

Although we showed the structure and activity of YjoB, several questions remain to understand the action mechanism of YjoB. First, conformational changes of YjoB by ATP hydrolysis should be further addressed. In our study, YjoB suppressed substrate aggregation in both the non-nucleotide- and AMPPNP-binding states, but not in the ADP-binding state. The nucleotide effect of this aggregation suppression pattern is similar to that of GroEL. This suggests that YjoB changes its conformation through ATP hydrolysis to release the bound substrate. However, it is unclear how YjoB binds and releases substrates through ATP hydrolysis. One possible model is the opening and closing of the internal chamber by the AAA domains of the YjoB hexamer (Supplementary Fig. S16). Unlike YjoB-NTD, which forms an interlocking hexamer, the YjoB-AAA domain can assemble and disassemble relatively freely, and its conformational changes by ATP hydrolysis can expose the internal chamber. In the crystal structure of ADP-bound YjoB, the electron density of ADP was alternately observed in three of the six AAA domains. A dimer unit of AAA domains containing a single ADP may open the internal chamber in the non-nucleotide or ATP-binding states and close the chamber when ADP binding occurs after ATP hydrolysis (Supplementary Fig. S16). However, no conformational changes between native YjoB and ADP-bound YjoB were observed in the crystal structures. As the nucleotide-dependent substrate binding of YjoB should be accompanied by conformational changes, the compact hexamer shown in the crystal structure of native YjoB may be caused by crystal packaging. Therefore, further studies are needed to investigate the reaction mechanism of ATP hydrolysis in YjoB. Second, the role of YjoB-binding proteins identified by AP-MS experiments should further address whether the proteins are substrates or adaptors for YjoB activity. Notably, SufA, YutI, and YjoA were highly ranked as YjoB-binding proteins in the AP-MS results. The proteins are thermally stable and small in size (less than 20 kDa). *yjoA* is located close to *yjoB* in the *B. subtilis* genome, but not in the same operon.

## METHODS

### Protein expression and purification

DNA encoding full-length YjoB (residues 1–423) was amplified from the genome of *B. subtilis* 168 strain using polymerase chain reaction and inserted into the pET-DUET1 vector (MilliporeSigma, St. Louis, MO, USA) modified for the N-terminal expression of 6×His and tobacco etching virus (TEV) protease cleavage site. The plasmid was then introduced into the *E. coli* strain BL21-star (DE3) (Thermo Fisher Scientific, Waltham, MA, USA), and the transformed cells were cultured in Luria–Bertani medium at 37 °C. Protein expression was induced by adding 0.4 mM isopropyl β-D-1-thiogalactopyranoside (IPTG) to the culture medium when the optical density at 600 nm (OD_600_) reached 0.6–0.7. After overnight culture at 20 °C, cells were harvested by centrifugation at 3,000 × *g* for 10 min and resuspended with buffer A (20 mM HEPES pH 7.5, 0.2 M NaCl, 0.2 mM TCEP, and 5% (v/v) glycerol). The resuspended cells were lysed by sonication and incubated with DNase I and RNase A at a concentration of 10 μg/mL for 30 min to digest the nucleic acids in the lysate. Insoluble debris was removed by centrifugation at 20,000 × *g* for 30 min.

YjoB was purified using IMAC and SEC. The clarified cell lysate was loaded onto a 5 mL HisTrap nickel chelating column (Cytiva, Marlborough, MA, USA). The resin was washed with 80 mM imidazole and proteins bound to the resin were eluted through an imidazole gradient of 0.08–1.0 M. Eluates containing 6×His-YjoB were treated with TEV protease at 4 °C. After the complete cleavage of 6×His-YjoB into 6×His and YjoB, the protein solution was dialyzed in buffer A for 3 h and passed through HisPur Ni-NTA resin (Thermo Fisher Scientific) to remove the 6×His tag. YjoB was further purified by SEC using a Superdex 200 preparatory grade column (Cytiva) equilibrated with buffer B (20 mM HEPES pH 7.5, 50 mM NaCl, 0.2 mM TCEP, and 5 % (v/v) glycerol). To prepare SeMet-labeled YjoB, BL21-star (DE3) cells transformed with the 6×His-YjoB plasmid were cultured in a minimal medium containing SeMet (TCI Chemicals, Tokyo, Japan). SeMet-labeled YjoB was purified using the same procedure as described for native YjoB.

To prepare the domains of YjoB, DNA encoding the NTD (residues 1–159) or the AAA domain (residues 159–423) of YjoB was inserted into the pET-DUET1 vector (MilliporeSigma) together with the DNA encoding the 6×His-SUMO-TEV protease cleavage site. The proteins were expressed and purified using the same procedure described for native YjoB. The YjoB-binding proteins, including AhpC, GapB, KatA, SufA, SufD, TrxA, YjoA, YphP, YqiW, YutI, and YwfI, were expressed and purified by the same procedures as YjoB domains. YjoB-NTD and KatA in different oligomer states were collected separately and further purified by SEC using a Superdex 200 preparatory column (Cytiva) equilibrated with buffer B.

To prepare GroEL, DNA encoding *B. subtilis* GroEL (residues 1-544) was inserted into the pET-22b vector (MilliporeSigma), and the plasmid was introduced in BL21-star (DE3) cells (Thermo Fisher Scientific). GroEL expression was induced by adding 0.4 mM IPTG to the culture medium when the OD_600_ reached 0.65. After culturing at 37 °C for 6 h, cells were harvested by centrifugation at 3,000 × *g* for 10 min and resuspended with buffer B. GroEL in cell lysates was precipitated with 60 % ammonium sulfate, collected by centrifugation at 10,000 × *g* for 30 min, and resuspended with buffer B. GroEL was further purified by SEC using a Superdex 200 preparatory grade column (Cytiva) equilibrated with buffer B. Molar concentrations of all proteins were calculated on a monomer basis.

### Crystallization, data collection, and structure determination

The crystals of YjoB and NTD-1 were obtained using the micro-batch method. Crystallization droplets were prepared by mixing 1 μL protein (YjoB and NTD-1 at 5 mg/mL) with 1 μL crystallization solution under Al’s oil layer in a micro-batch plate (Hampton Research, Aliso Viejo, CA, USA) and then incubated at 20 °C. YjoB crystals completely grew within 5 days in a crystallization solution containing 5% (v/v) PEG400, 2 M ammonium citrate/citric acid pH 7.5, and 10 mM TCEP. NTD-1 crystals grew within a week in a crystallization solution containing 30% (w/v) isopropanol, 0.1 M Tris-HCl (pH 8.5), and 30% (w/v) PEG3350. For crystallization of ADP-bound YjoB, YjoB at 10 mg/mL was mixed at a 1:1 volume ratio with buffer C (20 mM HEPES at pH 7.5 with 50 mM NaCl, 0.2 mM TCEP, 5% (v/v) glycerol, 1 mM ADP, and 2 mM MgCl_2_), and YjoB-ADP crystals were obtained using the same methods used to obtain YjoB crystals.

Diffraction data of crystals were collected at PLSII-BL7A and 11C (Beamline 7A and 11C, Pohang Light Source II, South Korea). For cryo-protection, 0.5 μL of 100% glycerol was added directly to the crystallization drop as a cryo-protectant. The crystal was then picked up using a crystal harvesting loop (MiTeGen, Ithaca, NY, USA) and flash-frozen in a cold nitrogen stream of the beamline hutch. Diffraction data were indexed, integrated, and scaled using MOSFLM (40) and HKL2000 (41).

The initial experimental electron density map of SeMet-labeled YjoB was obtained through the MAD phasing method. The structure model was built by tracing the electron density and improved by iterating structure refinement and model building. The crystal structures of native YjoB, YjoB-ADP, and YjoB-NTD were determined via the molecular replacement phasing method using the structural model of SeMet-labeled YjoB as a template. R/R_free_ values of the final structures for native YjoB, YjoB-ADP, and YjoB-NTD were 20.0/23.6, 18.7/22.3, and 23.7/28.0 (%), respectively. MAD phasing, molecular replacement phasing, model building, and structure refinement were performed using PHENIX.autosol (42), Phaser (43), COOT (44), and PHENIX.refine (45) programs, respectively. Data collection and refinement statistics are summarized in Supplementary Table S1. The figures were drawn using PyMOL (46) and ALSCRIPT (47). Surface area, surface charge, protein-protein interaction, and structural alignment were analyzed using PISA (25), APBS (48), DIMPLOT (49), and the DALI server (24), respectively.

### Affinity purification and mass spectroscopy

DNAs encoding 3×FLAG-2×Strep-YjoB and 3×FLAG-2×Strep-EGFP were inserted into the shuttle vector pHT254 (MoBiTec, Göttingen, Germany). Each plasmid was introduced into *B. subtilis* 168 strain (ATCC-23857, ATCC, Manassas, VA, USA), and the transformed *B. subtilis* cells were cultured in 100 mL Lennox Luria–Bertani medium at 37 °C. When the OD_600_ of the medium reached 0.6–0.7, the protein expression was induced by adding 0.5 mM IPTG into the culture medium. After overnight culture at 20 °C, the cells were harvested by centrifugation at 3,000 × *g* for 10 min, resuspended in PBS (4.3 mM Na_2_HPO_4_, 1.4 mM KH_2_HPO_4_, 137 mM NaCl, and 2.7 mM KCl) supplemented with a protease inhibitor cocktail (Thermo Fischer Scientific), and lysed by sonication. Cell lysates were incubated with DNase I and RNase A at 10 μg/mL for 30 min and clarified by centrifugation at 20,000 × *g* for 30 min.

3×FLAG-2×Strep-YjoB and 3×FLAG-2×Strep-EGFP were purified using Strep-pulldown and FLAG-IP. Cell lysates (5 mL) were incubated with 100 μL pre-clearing beads (Thermo Fischer Scientific) for 1 h, followed by centrifugation at 1000 × *g* for 2 min to remove the beads. The pre-cleared cell lysates were mixed with 150 μL Strep-Tactin Sepharose (IBA, Göttingen, Germany). After 3 h incubation at 4 °C, the Strep resin was washed three times with PBS containing 0.05% Nonidet P-40. Proteins bound to the resin were eluted by incubation with 240 μL E buffer (IBA) for 30 min followed by centrifugation at 1000 × *g* for 1 min. For the second affinity purification, the eluates from Strep-pulldown were diluted 2-fold with PBS buffer and incubated with 60 μL anti-FLAG M2 affinity gel (MilliporeSigma) for 1 h. The resin was then washed three times with PBS containing 0.05% NP40. Proteins bound to the resin were eluted by incubation with 100 μL PBS supplemented with 0.05% NP40 and 500 μg/mL 3×FLAG peptide for 30 min followed by centrifugation at 1000 × *g* for 1 min. The buffer was exchanged with 50 mM ammonium bicarbonate using an Amicon Ultra centrifugal filter with a 10 kDa molecular weight cutoff (MilliporeSigma). The protein concentration of the eluate was then quantified using a BCA assay kit (Thermo Fischer Scientific).

For the LC-MS/MS analysis, 7 ug of protein was denatured at 95 °C for 5 min, reduced with 10mM DTT at 60 °C for 1 h, and alkylated with 20 mM iodoacetamide for 40 min in the dark. Protein samples were mixed with Trypsin Gold (Promega, Madison, WI, USA) at a 50:1 ratio (w/w) and incubated at 37 °C for 16 h. The digested protein sample was desalted with a C18 tip (Thermo Fisher Scientific), dried in a speed vacuum concentrator, and dissolved in 50 mL water containing 0.1% formic acid. The sample was then injected into an Acclaim PepMap100 RSLC C18 column (75 μm x 50 cm, 2 μm bead size, 100 Å pores) connected to an Ultimate 3000 RSLCnano system with an Orbitrap Eclipse Tribrid mass spectrometer (Thermo Fisher Scientific). Three independent affinity purifications and MS/MS analyses were performed each for YjoB and EGFP.

Mass spectrometric raw data were analyzed using the MaxQuant Software Package v2.0.3.1 (50). The derived peak list was searched against the *B. subtilis* 168 strain proteome (Uniprot taxon identifier 224308) and EGFP sequences using the database search engine Andromeda(51). In the analysis, trypsin/P was specified as the enzyme allowing up to two missing cleavages. Carbamidomethylation on Cys was specified as a fixed modification, whereas oxidation on Met was specified as a variable modification. Label-free quantification (LFQ) option was activated with a minimum ratio count of 2. All the other parameters in MaxQuant were set to default values. The MaxQuant output tables showing the lists of identified proteins were statistically analyzed using the Perseus software suite v2.0.3.0(52). Common contaminants, reverse hits, and identification by site were removed from the tables. LFQ values were transformed to the logarithm (log_2_) and divided into two groups (YjoB and EGFP). Proteins identified with at least two valid values from three LFQ values in a single group were selected, and the missing values were imputed by a normal distribution with a downshift of 1.2 and a width of 0.30. Statistical significance between YjoB and EGFP groups was calculated with a two-tailed t-test. The *p*-value (-log_10_) and LFQ intensity difference between YjoB and EGFP groups were plotted using SigmaPlot 14.0 (Systat Software, San Jose, CA, USA). emPAI values of identified proteins were calculated using the Integrated Proteomics Pipeline v5.0.1 (Integrated Proteomics Application, San Diego, Ca, USA).

### SEC-MALS

Protein molar mass was measured using a SEC-MALS instrument (Wyatt Technology, Santa Barbara, CA, USA). A volume of 100 μL YjoB (5 mg/mL) was injected into a Superdex200 Increase 10/300 GL analytical column equilibrated with phosphate-buffered saline (10 mM Na_2_HPO_4_, 1.8 mM KH_2_PO_4_, 2.7 mM KCl, and 0.137 M NaCl; pH 7.4), and the eluate was applied to inline DAWN Heleos II MALS and Optilab T-Rex differential refractive index detectors (Wyatt Technology). Data were analyzed using the ASTRA 6 software package (Wyatt Technology) and visualized using SigmaPlot 14.0 (Systat Software, San Jose, CA, USA).

### ATPase activity measurement

ATPase activity was measured by observing a decrease in NADH levels with an increase in ADP levels (53). For enzyme kinetics analysis, 100 μL reaction mixtures containing 100 nM YjoB, 1 – 1,000 μM ATP, 1 μL pyruvate kinase/lactic dehydrogenase enzymes from rabbit muscle (MilliporeSigma), 4 mM phosphoenolpyruvate, 4 mM MgCl_2_, 150 mM KCl, and 0.32 mM NADH in buffer A were placed onto a 96-well microtiter plate (Corning Incorporated, Corning, NY, USA). NADH oxidation was monitored at 340 nm with 1 min intervals for 30 min at 30 °C using a SpectraMax iD5 microplate reader (Molecular Devices, San Jose, CA, USA).

### Chaperone activity measurement

Chaperone activity was measured by observing the thermal aggregation of proteins (54). Mixtures (200 μL) containing YjoB and target proteins in buffer A were placed onto a 96-well microtiter plate (Corning Incorporated). Absorbance at 340 nm was recorded for 150 min with 3 min intervals using a SpectraMax iD5 microplate reader (Molecular Devices) preheated to 42 °C. The mean ± standard error of three replicates was analyzed and visualized using SigmaPlot 14.0 (Systat Software).

### Catalase activity measurement

Catalase activity was assessed by comparing the H_2_O_2_ levels that remained after incubation with KatA. KatA was purified in three different oligomeric states (monomer: KatA-1; dimer: KatA-2; and larger oligomer: KatA-3). Mixtures (100 μL) containing 0.5 μM YjoB, 0.5 μM KatA, and 100 μM ATP in buffer A were incubated at 30 °C or 42 °C for 1 h, followed by further incubation with hemin at 30 °C for 30 min. H_2_O_2_ was added to the pre-incubated mixture, and 200 μL reaction mixtures were placed onto 96-well microtiter plates (Greiner Bio-one, Kremsmünster, Austria). Absorbance at 240 nm was recorded (55) at 30 °C for 20 min using a SpectraMax iD5 microplate reader (Molecular Devices). The reaction mixture contained 25 nM KatA-1 (or 0.25 nM KatA-2 or 0.25 nM KatA-3), 25 nM YjoB, 5 μM ATP, 25 nM hemin, and 20 mM H_2_O_2_. The mean ± standard error of three replicates was analyzed and visualized using SigmaPlot 14.0 (Systat Software).

To compare the catalase activity of wild-type *B. subtilis* and its mutants (*ΔkatA* and *ΔyjoB*), cell lysates were prepared. Wild-type *B. subtilis* and its mutants were cultured in 100 mL Lennox Luria–Bertani media at 37 °C and harvested at exponential (OD_600_ = 0.65), early stationary (OD_600_ = 1.0), and extended stationary (2 h after reaching OD_600_ = 1.0) phases. Cells in a 15 mL medium were harvested by centrifugation at 3,000 × *g* for 10 min. Collected cells were resuspended in buffer A containing a protease inhibitor cocktail (Thermo Fisher Scientific), lysed by sonication, and clarified by centrifugation at 20,000 × *g* for 10 min at 4 °C. The protein concentration of the lysate was measured using a BCA assay kit (Thermo Fisher Scientific). Reaction mixtures (200 μL) containing 1.0 μg protein and 20 mM H_2_O_2_ in buffer A were placed onto 96-well microtiter plates (Greiner Bio-one). H_2_O_2_ decomposition was measured at 30 °C for 20 min. Three *B. subtilis* strains (56) (*B. subtilis* subsp. *subtilis* (BGSC ID: 1A1), Δ*katA::kan* (BGSC ID: BKK08820), and Δ*yjoB::kan* (BGSC ID: BKK12420)) were obtained through the Bacillus Genetic Stock Center (Columbus, OH, USA).

To measure the catalase activity of renatured KatA-1, 40 μM KatA-1 was denatured with 6 M guanidine hydrochloride at 37 °C for 1 h and then renatured by diluting 80-fold in buffer A containing YjoB and hemin. The concentrations of KatA-1, YjoB, and hemin in the diluted buffer were 500 nM. Catalase activity was measured with 200 μL reaction mixtures containing 25 nM KatA-1, 25 nM YjoB, 20 mM H_2_O_2_, and 25 nM hemin in buffer A.

## ACKNOWLEDGMENTS

We would like to thank the staffs (Dr. Geul Bang and Dr. Eunha Hwang) of the Korea Basic Science Institute (KBSI) for their assistance in mass spectrometry and SEC-MALS analyses. This research was supported by the Basic Science Research Program through the National Research Foundation of Korea (NRF) funded by the Ministry of Education [grant numbers NRF-2020R1I1A1A01060732 and NRF-2020R1I1A3060361].

## DATA AND CODE AVAILABILITY

The final coordinates and structure factors that support the findings of this study have been deposited in the Protein Data Bank with the PDB accession numbers [7W42 for YjoB, 7W46 for YjoB-ADP and 7W43 for YjoB-NTD]. The mass spectrometry proteomics data have been deposited to the ProteomeXchange Consortium via the PRIDE(57) partner repository with the dataset identifier PXD032982 and 10.6019/PXD032982. Source data are provided with this paper.

## AUTHOR CONTRIBUTIONS

**Eunju Kwon:** Conceptualization, Methodology, Validation, Formal analysis, Investigation, Resources, Data curation, Writing-Original draft, Visualization, Project administration, Funding acquisition. **Pawan Dahal:** Conceptualization, Methodology, Formal analysis, Investigation, Resources, Writing-Original draft. **Dong Young Kim:** Conceptualization, Writing-Reviewing and Editing, Visualization, Supervision, Funding acquisition.

