## Supplementary Figures and Tables for "Crystal structure and biochemical analysis suggest that YjoB ATPase is a substrate-specific molecular chaperone"

**Supplementary Figure S1**

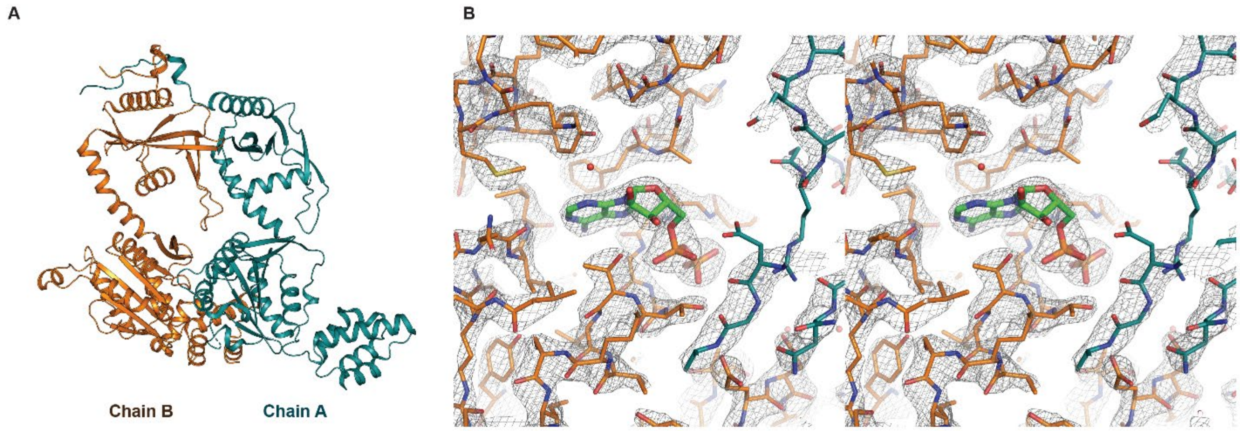

**Supplementary Figure S1. Two monomers in the asymmetric unit of the YjoB crystal.** (**A**) The two chains of the asymmetric unit are drawn as teal and orange ribbon models. (**B**) The 2Fo-Fc electron density map of YjoB around the ADP-binding site is displayed in stereo view at a contour level of 2.0.

### Supplementary Figure S2

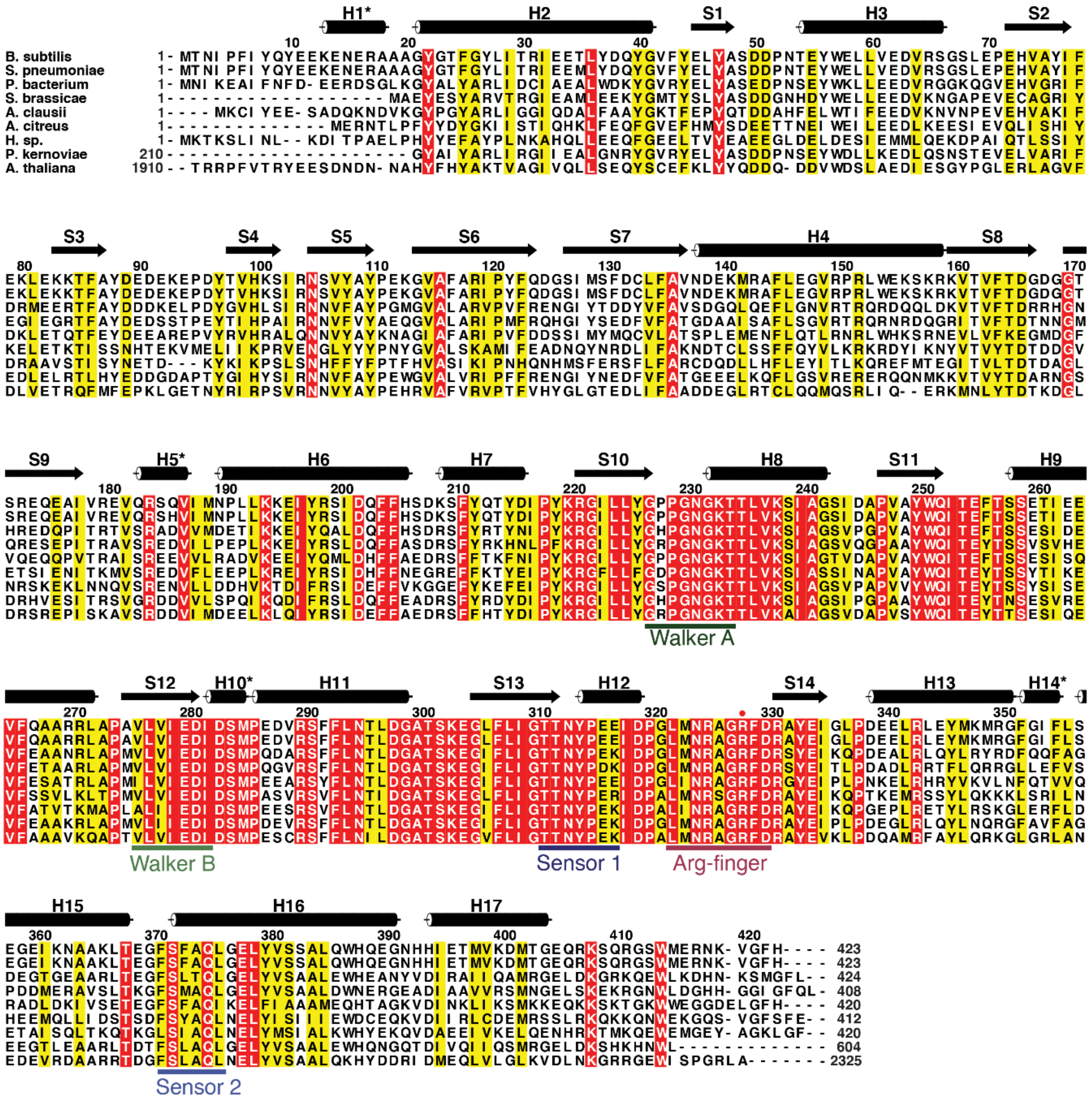

**Supplementary Figure S2. Sequence alignment of YjoB homologs.** Cylinders and arrows represent helices and strands of *B. subtilis* YjoB, respectively. Asterisk (*) indicates the $3^{10}$-helix. YjoB homologs and their abbreviations used in the sequence alignment are as follows: *Streptococcus pneumoniae* cell-division protein (GenBank ID: CJR51781.1): *S. pneumoniae*; *Paenibacillaceae bacterium* ATP-binding protein (GenBank ID: RXZ82979.1): *P. bacterium*; *Saccharibacillus brassicae* ATP-binding protein (GenBank ID: QDH23826.1): *S. brassicae*; *Alkalihalobacillus clausii* ATPase (GenBank ID: AST95141.1): *A. clausii*; *Arthrobacter citreus* ATP-binding protein (GenBank ID: QKE75788.1): *A. citreus*; *Halomonas* sp. MG34 AAA family ATPase (GenBank ID: NAO99206.1): H. sp; *Phytophthora kernoviae* 00238/432 hypothetical protein G195_000577 (GenBank ID: KAF4325776.1): *P. kernoviae*; and *Arabidopsis thaliana* hypothetical protein AXX17_ATUG04490 (GenBank ID: OAO89099.1): *A. thaliana*.

### Supplementary Figure S3

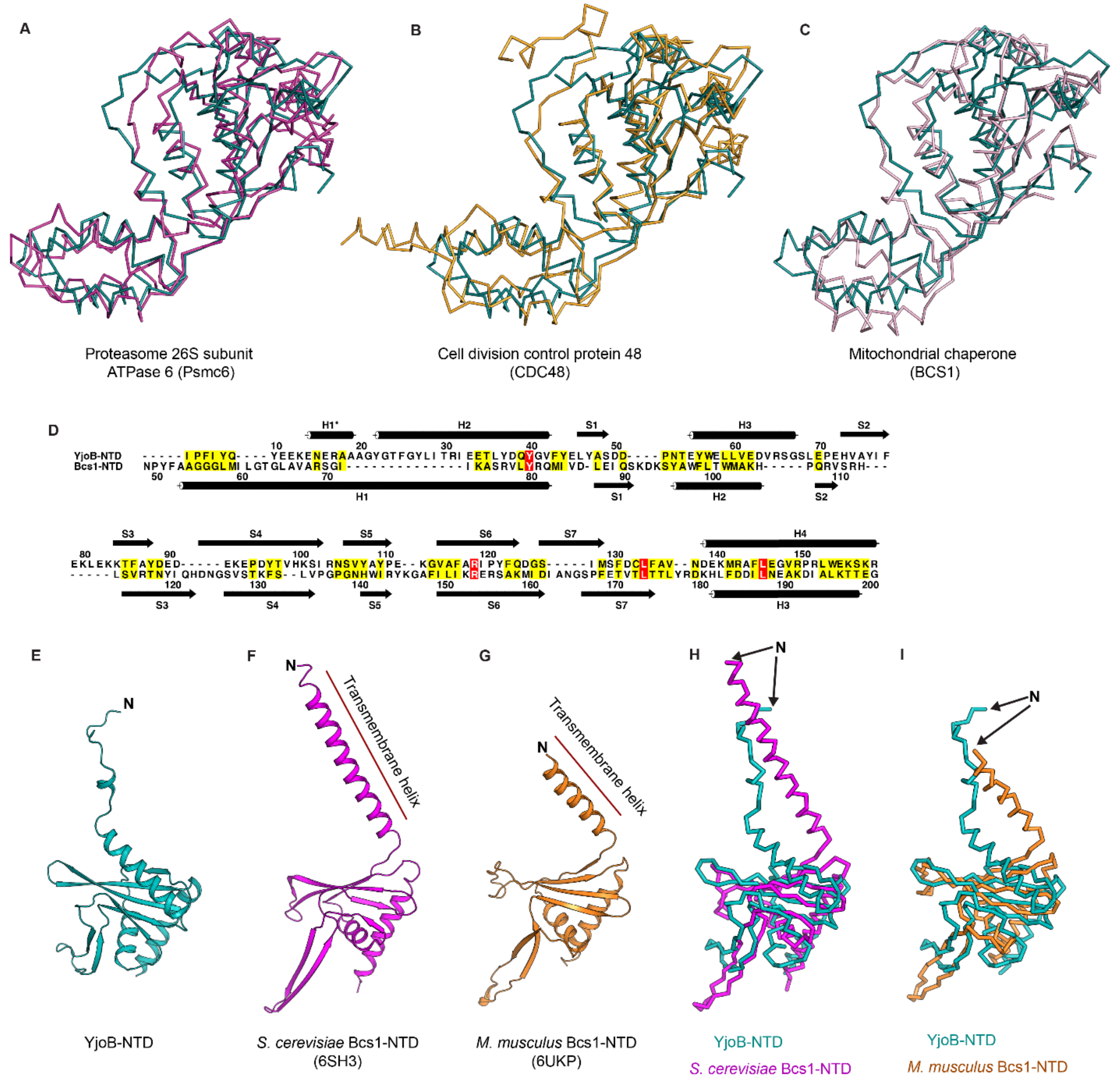

**Supplementary Figure S3. Structural homologs of the NTD and AAA domain of YjoB.** Cα trace models of the AAA domains superimposed onto the YjoB-AAA domain (deep teal). (**A**) *Rattus norvegicus* Psmc6 (PDB ID: 6EPC). (**B**) *Saccharomyces cerevisiae* CDC48 (PDB ID: 6OPC). (**C**) *Saccharomyces cerevisiae* Bcs1 (PDB ID: 6SH3). The sequence identities of the AAA domains of Psmc6, CDC48, and Bcs1 aligned with the YjoB-AAA domain are 30.7, 29.6, and 32.8 (%), respectively. (**D**) Structure-based sequence alignment of YjoB-NTD and *S. cerevisiae* Bcs1-NTD. Cylinders and arrows represent helices and strands. (**E-G**) Ribbon models of YjoB-NTD (E), *S. cerevisiae* Bcs1-NTD (PDB ID: 6SH3) (F), and *Mus musculus* Bcs1-NTD (PDB ID: 6UKP) (G) shown in the same orientation. (**H, I**) Cα trace models of *S. cerevisiae* Bcs1-NTD (H) and *M. musculus* Bcs1-NTD (I) superimposed onto the YjoB-NTD.

### Supplementary Figure S4

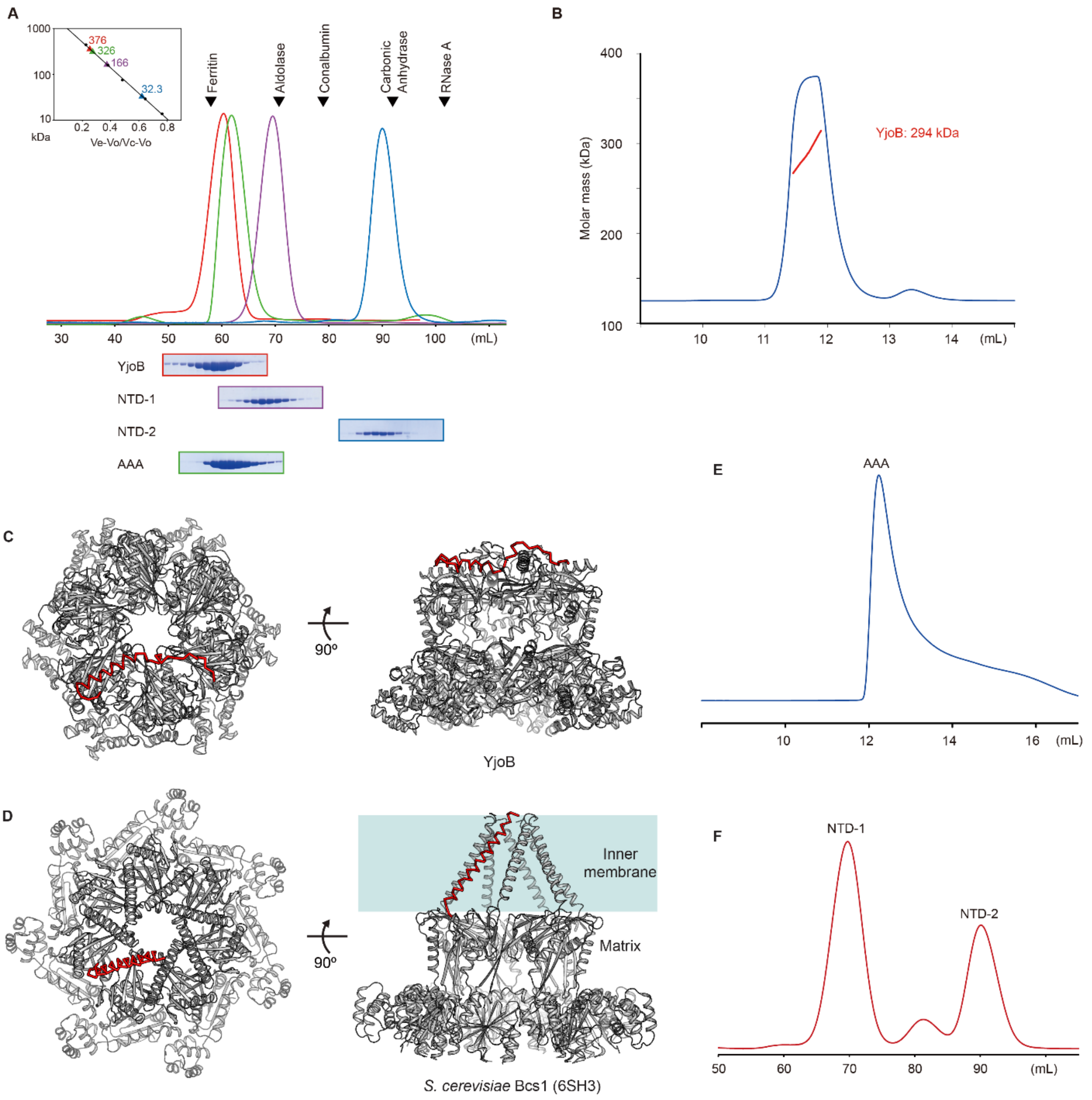

**Supplementary Figure S4. YjoB oligomerization.** (**A**) SEC of YjoB, NTD-1, NTD-2, and YjoB-AAA domain. UV profiles of YjoB, NTD-1, NTD-2, and YjoB-AAA domains from the Superdex200 preparatory column are shown in red, violet, blue, and green, respectively. Molar mass estimated from molecular standards is shown in the left box. Ferritin (440 kDa), aldolase (158 kDa), conalbumin (75 kDa), carbonic anhydrase (29 kDa), and RNase A (13.7 kDa) were used as molecular standards. The eluate of each UV peak is shown in the SDS-PAGE. (**B**) SEC-MALS of YjoB. The red line represents the molar mass of the eluate (approximately 294 kDa). YjoB is estimated to be a hexamer. (**C**) Ribbon models of the YjoB hexamer shown in top and side views. The N-terminus containing helices H1 and H2 is represented by the red Cα trace model. (**D**) Ribbon models of *S. cerevisiae* Bcs1 (PDB ID: 6SH3) shown in top and side views. The N-terminal transmembrane helix is represented by the red Cα trace model. (**E**) SEC profile of the YjoB-AAA domain showing that the AAA domain is heterogeneous in the PBS. (**F**) SEC result of the YjoB-NTD during the purification. The YjoB-NTD was separated into a monomer and a larger oligomer using a Superdex200 preparatory column.

**Supplementary Figure S5**

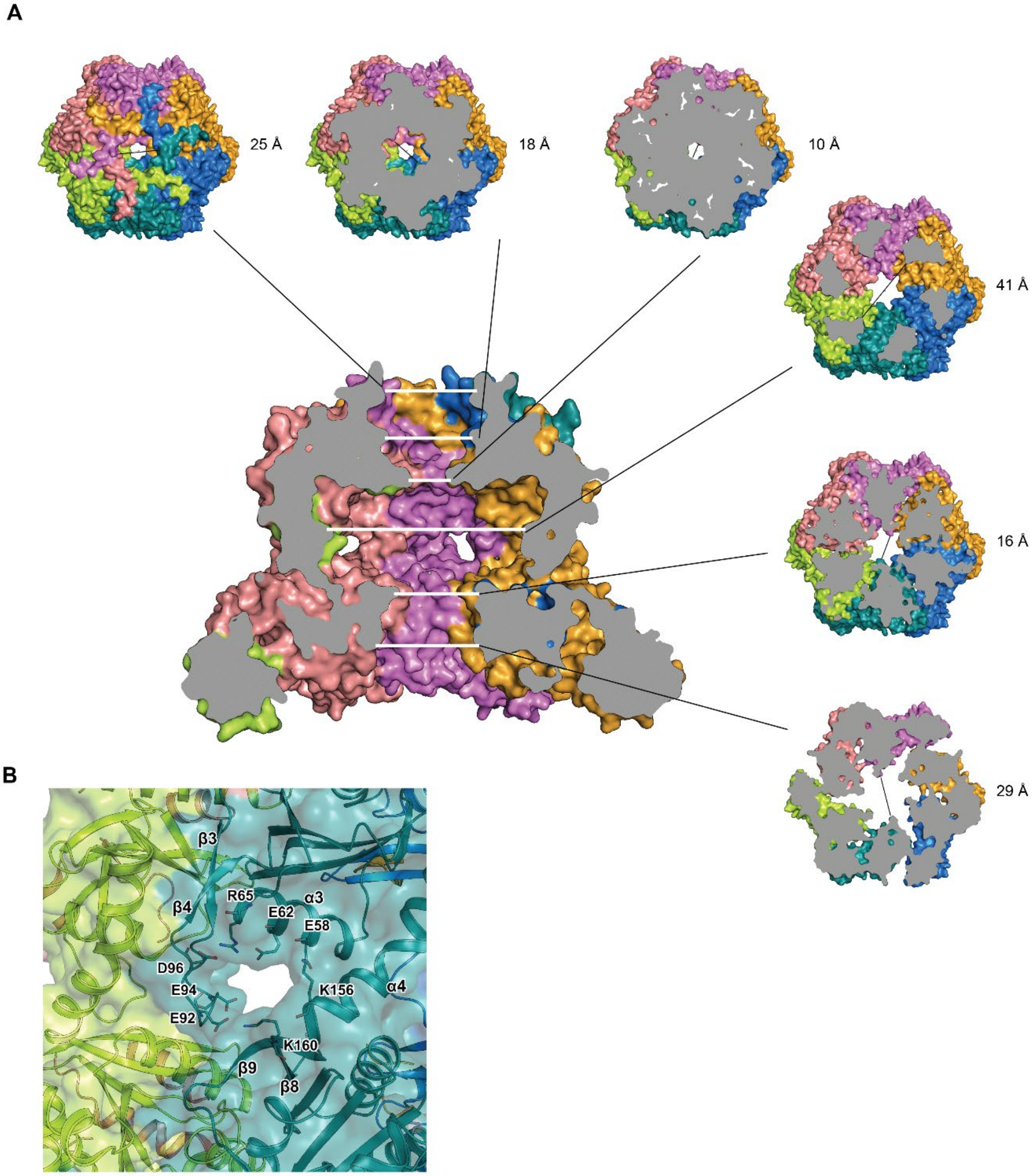

**Supplementary Figure S5. Central and lateral pores of the YjoB hexamer.** (**A**) Cross-sectional surface model of the side view. The central pore diameter is shown as a cross-sectional model in a top view. (**B**) Transparent surface model showing a lateral pore. The residues involved in the formation of the pore are shown as stick models and labeled.

**Supplementary Figure S6**

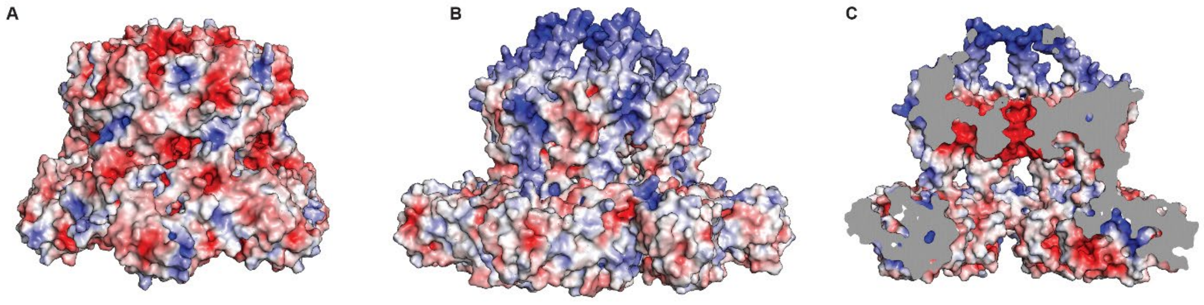

**Supplementary Figure S6. Surface charge distribution models of YjoB and Bcs1. Surface charge distribution models of YjoB and Bcs1.** (**A**) Charge distribution on external surfaces of the YjoB hexamer. (**B, C**) Charge distribution on external (**B**) and internal surfaces (**C**) of the Bcs1 heptamer (PDB ID: 6SH3). Electrostatic potential from red (−5 kT/e) to blue (+5 kT/e) is plotted on the solvent-accessible surface.

**Supplementary Figure S7**

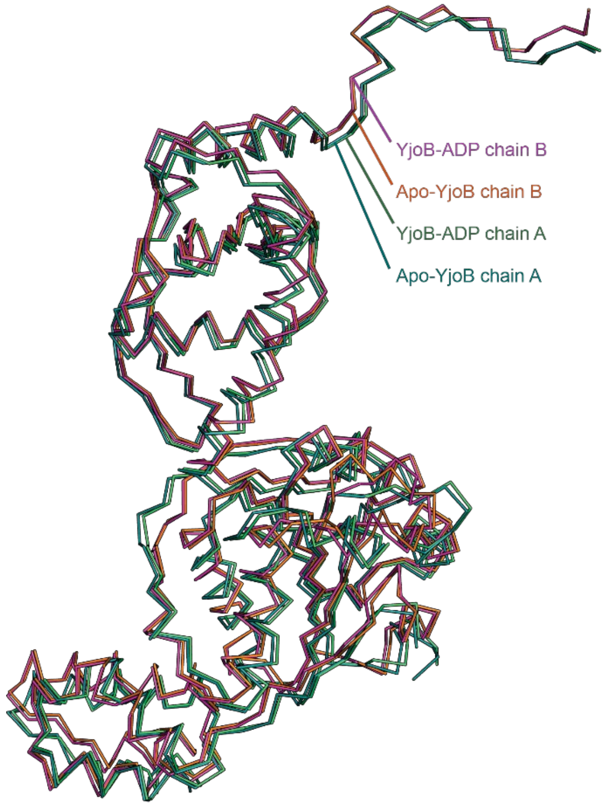

**Supplementary Figure S7. Structure comparison of YjoB and YjoB-ADP.** Cα trace models of the YjoB monomers. Monomers in the asymmetric unit of the YjoB and YjoB-ADP crystals are superimposed. Chains A and B of YjoB and chains A and B of YjoB-ADP are in deep teal, green, red, and violet, respectively.

### Supplementary Figure S8

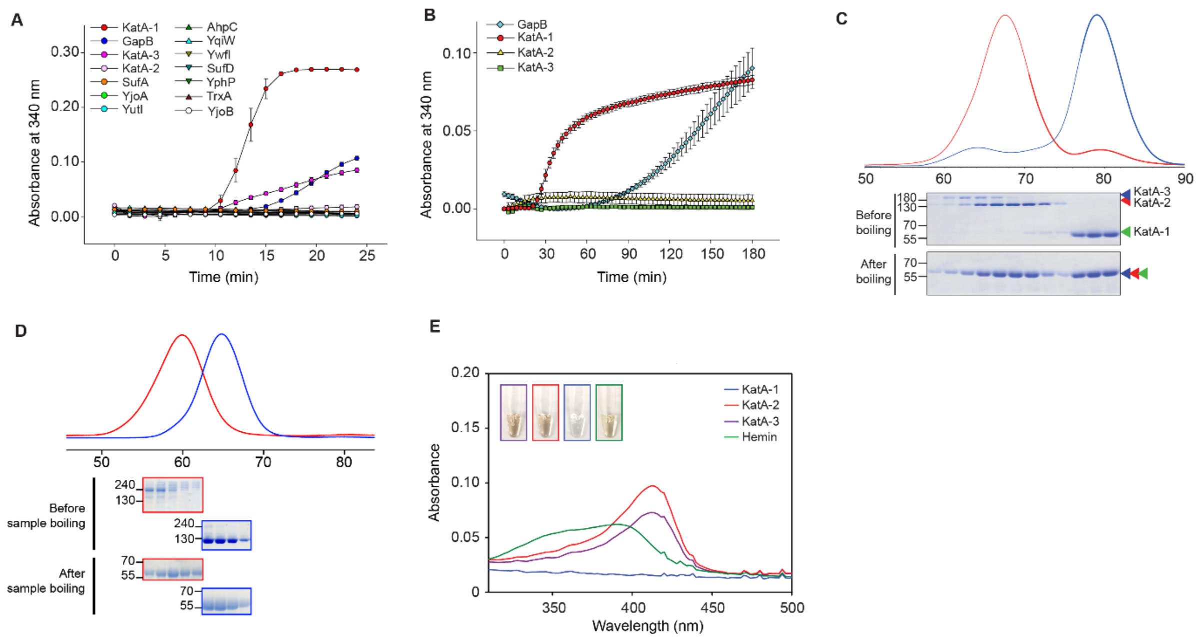

**Supplementary Figure S8. Thermal aggregation of YjoB-binding proteins.** (**A**) Thermal aggregation of YjoB and its binding proteins. Aggregation levels of AhpC, GapB, SufA, SufD, KatA-1, KatA-2, KatA-3, YjoA, YjoB, YphP, YqiW, YutI, and YwfI (10 μM each) were observed at 64 °C for 25 min. KatA and GapB form aggregates, whereas others are heat-stable. (**B**) Aggregation of KatA (5 μM) and GapB (10 μM) at 42 °C. KatA-1 and GapB form aggregates, whereas KatA-2 and KatA-3 are stable at 64 °C. Data in (A) and (B) are presented as the mean ± standard error of three replicates. (**C**) SEC results of KatA during purification. KatA is separated into a monomer (KatA-1), a dimer (KatA-2), and a higher oligomer (KatA-3). (**D**) SEC results of purified KatA-2 and KatA-3. KatA-2 and KatA-3 were purified without cross-contamination. (**E**) Absorption spectra of hemin bound to KatA. Absorbance (300 to 500 nm) of KatA oligomers (1 μM) and hemin (5 μM) was measured at 20 °C. KatA-2 and KatA-3 are shown in brown, with an absorption peak at 412 nm, whereas KatA-1 is shown colorless, with no absorption peak, indicating that KatA-1 is an inactive form that does not contain heme.

### Supplementary Figure S9

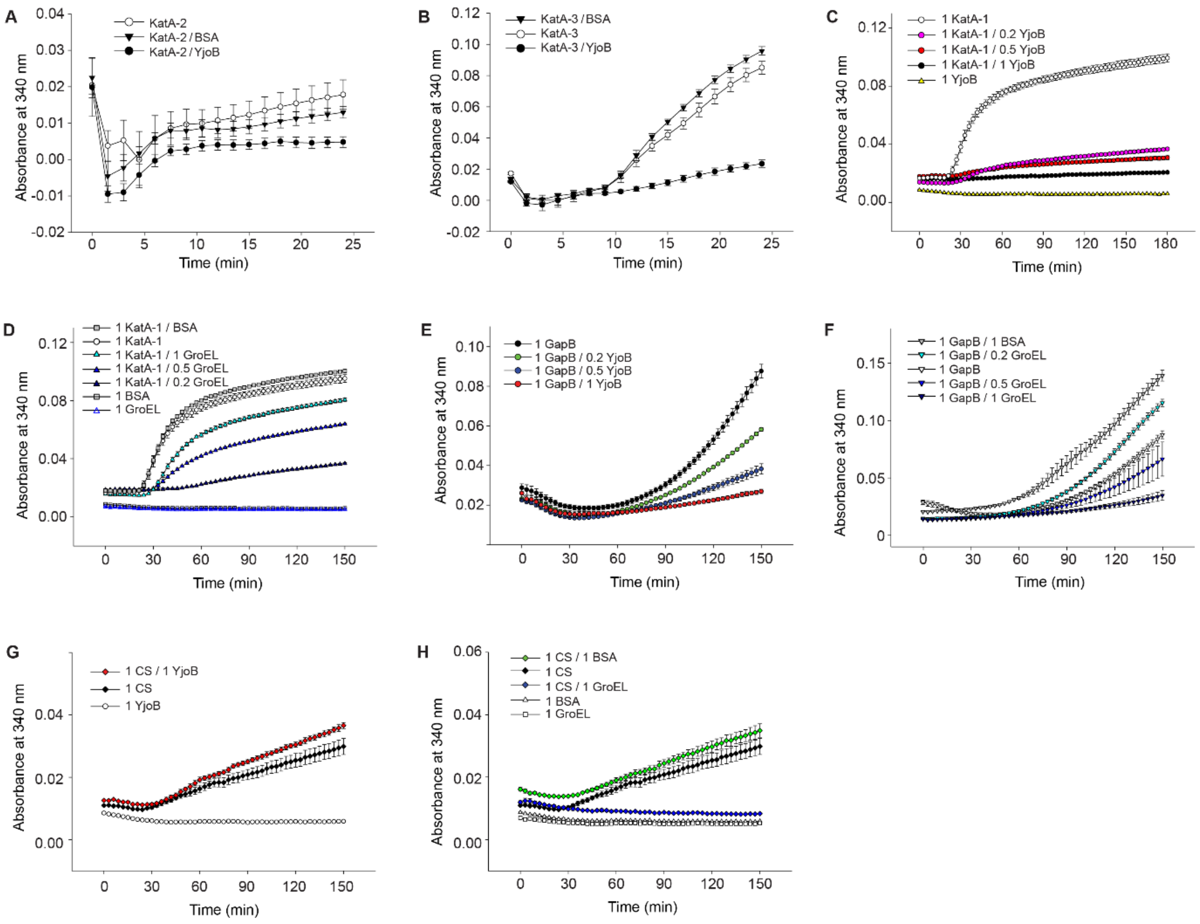

**Supplementary Figure S9. Chaperone activity of YjoB.** (**A, B**) Absorbance curves showing thermal aggregation of KatA-2 (A) and KatA-3 (B). KatA-2 and KatA-3 (10 µM each) were mixed with 5 µM YjoB, and the level of aggregation was observed at 64 °C for 25 min. (**C, D**) Thermal aggregation of KatA-1 (5 µM) in the presence of YjoB (C), GroEL, or BSA (D). (**E, F**) Thermal aggregation of GapB (10 µM) in the presence of YjoB (E), GroEL, or BSA (F). (**G, H**) Thermal aggregation of CS (10 µM) in the presence of YjoB (G), GroEL, or BSA (H). (**C–H**) 5 µM KatA-1, 10 µM GapB, and 10 µM CS were used as substrates. Molar ratios between proteins (monomer basis) are labeled in each panel. Absorbance curves at 340 nm are recoded at 42 °C for 150 min. These spectra show that the aggregation of KatA-1 and GapB is suppressed by both YjoB and GroEL, but not by BSA. CS aggregation was suppressed only by GroEL. Data are presented as the mean ± standard error of three replicates. Aggregation levels at the 150 min time point of (C)–(H) are shown as bar diagrams in Fig. 5, A to C.

### Supplementary Figure S10

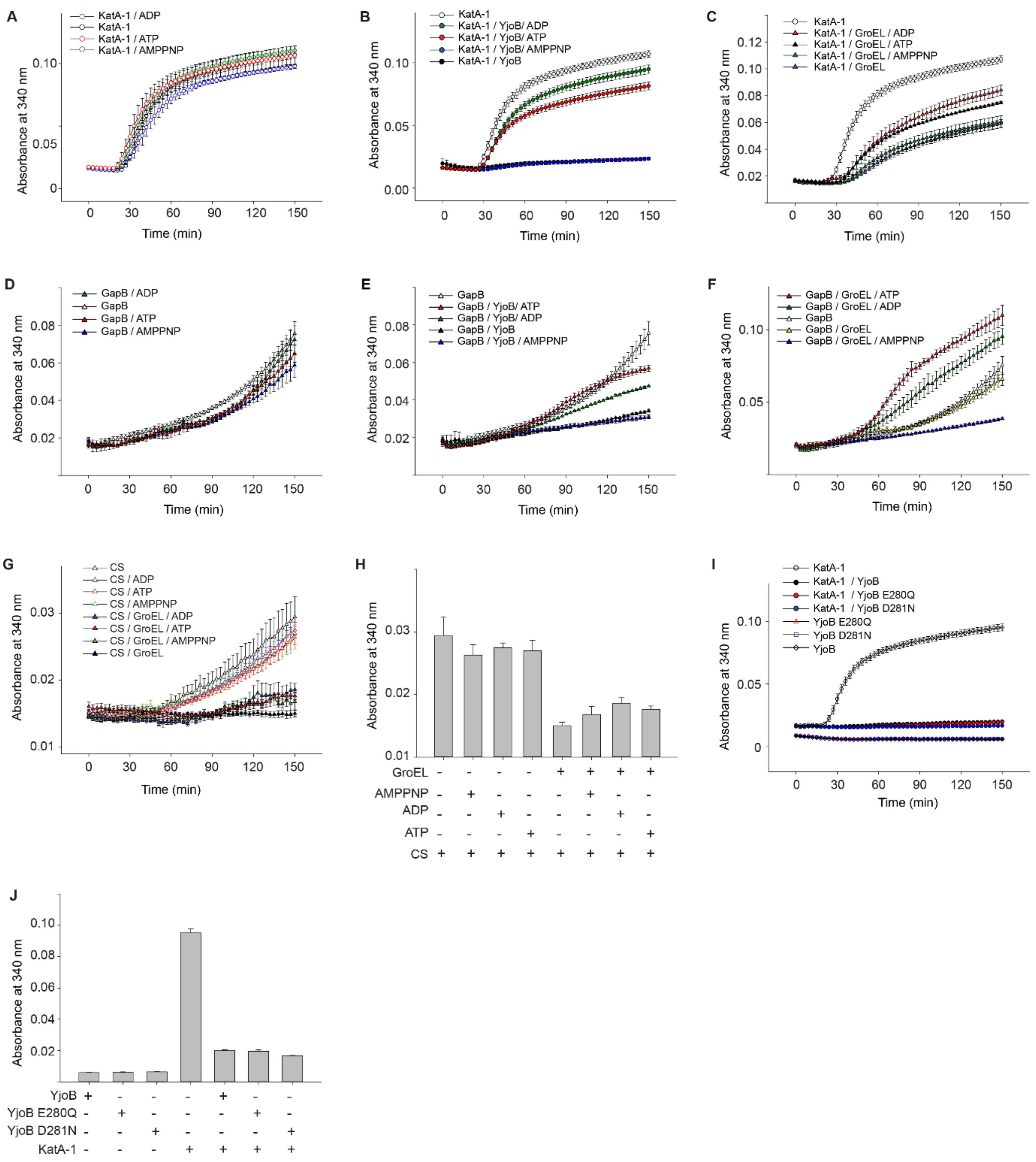

**Supplementary Figure S10. Chaperone activity of YjoB in the presence of adenine nucleotides.** (**A–C**) Nucleotide effects on the thermal aggregation of KatA-1. The reaction mixture contained 5 μM KatA-1, 2.5 μM GroEL, 2.5 μM YjoB, and 100 μM adenine nucleotide (ADP, ATP, or AMPPNP). (**A**) Thermal aggregation of KatA-1 in the presence of nucleotides. (**B**) Thermal aggregation of KatA-1 in the presence of YjoB and nucleotides. (**C**) Thermal aggregation of KatA-

1 in the presence of GroEL and nucleotides. (**D–F**) Nucleotide effects on the thermal aggregation of GapB. The reaction mixture contained 10 μM GapB, 5 μM GroEL, 5 μM YjoB, and 100 μM adenine nucleotide (ADP, ATP, or AMPPNP). (**D**) Thermal aggregation of GapB in the presence of nucleotides. (**E**) Thermal aggregation of GapB in the presence of YjoB and nucleotides. (**F**) Thermal aggregation of GapB in the presence of GroEL and nucleotides. (**G**) Nucleotide effects on the thermal aggregation of CS. The reaction mixture contained 10 μM CS, 5 μM GroEL, and 100 μM adenine nucleotides. (**H**) Bar diagrams showing the absorbance at the 150 min time point in the absorbance curves in (G). (**I**) Thermal aggregation of KatA-1 in the presence of YjoB mutants (E280Q and D281N). KatA-1 (5 μM) was mixed with YjoB or YjoB mutants at a 1:1 ratio. (**J**) Bar diagrams show absorbance at the 150 min time point of (I). All experiments were performed at 42 °C. Data are presented as the mean ± standard error of three replicates.

### Supplementary Figure S11

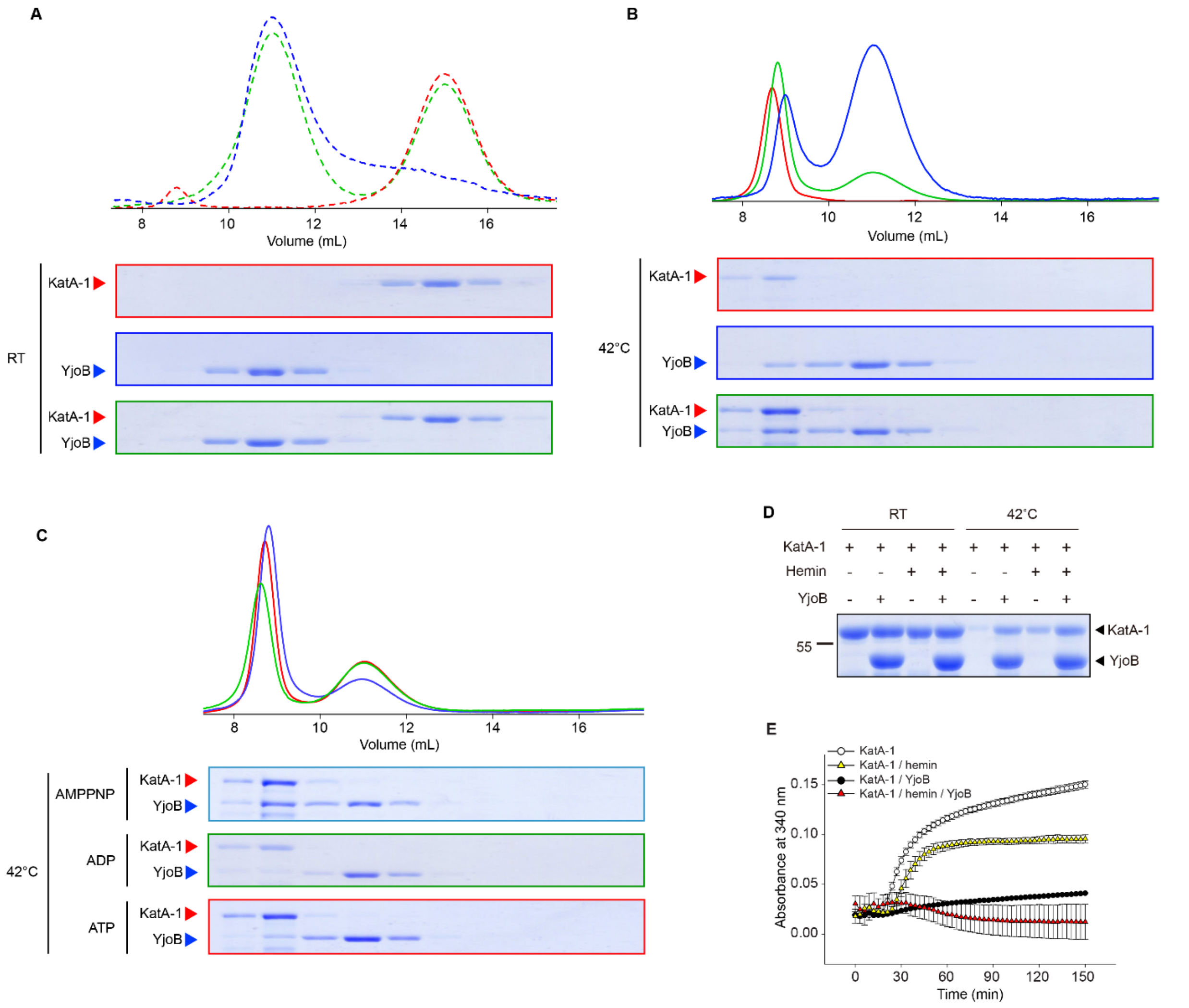

**Supplementary Figure S11. Interaction between YjoB and KatA-1.** (**A, B**) Interactions between YjoB and KatA-1. The KatA-1/YjoB mixtures were pre-incubated at 20 °C. (A) and 42 °C (B) for 1 h and injected onto an analytical Superdex200 column. KatA-1 monomer does not interact with YjoB at 20 °C (A), whereas the heat-induced oligomer of KatA-1 interacts with YjoB. (**C**) Interaction between YjoB and heat-induced KatA-1 oligomer in the presence of AMPPNP, ADP, or ATP. The mixture was pre-incubated at 42 °C for 1 h and injected onto an analytical Superdex200 column. KatA-1 co-eluted with YjoB in the presence of AMPPNP but not in the presence of ADP or ATP. (**D**) SDS-PAGE of YjoB and KatA-1. The YjoB/KatA-1 mixture was incubated at room temperature and 42 °C for 1 h, and their soluble fractions were collected after centrifugation. KatA-1 stability at 42 °C is partially increased by hemin and further increased by YjoB. (**E**) Hemin effects on the thermal aggregation of KatA-1. The reaction mixture contained 5 μM KatA-1, 5 μM YjoB, and 5 μM hemin. The experiments were performed at 42 °C, and the data are presented as the mean ± standard error of three replicates.

**Supplementary Figure S12**

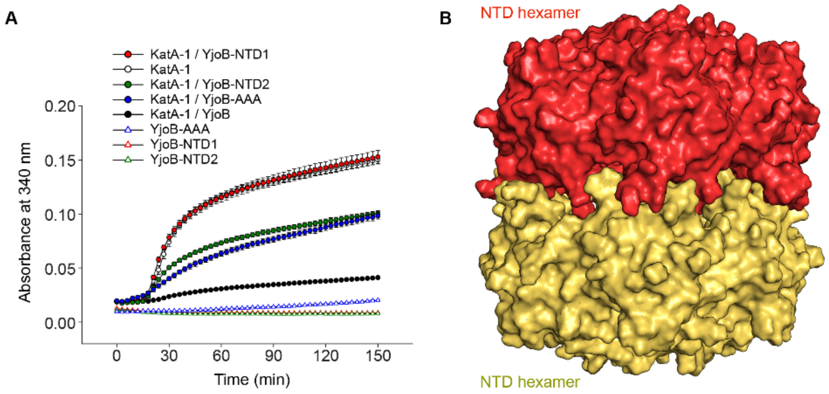

**Supplementary Figure S12. Role of the internal chamber in YjoB chaperone activity.** (**A**) Thermal aggregation of KatA-1 in the presence of YjoB domains. YjoB-NTD was purified as an oligomer (NTD-1) and a monomer (NTD-2). The aggregation of KatA-1 (10 μM) was compared in the presence of YjoB, NTD-1, NTD-2, and YjoB-AAA (5 μM each). Absorbance was measured at 42 ˚C for 150 nm, and the data are presented as the mean ± standard error of three replicates. (**B**) Surface model of the NTD-1.

**Supplementary Figure S13**

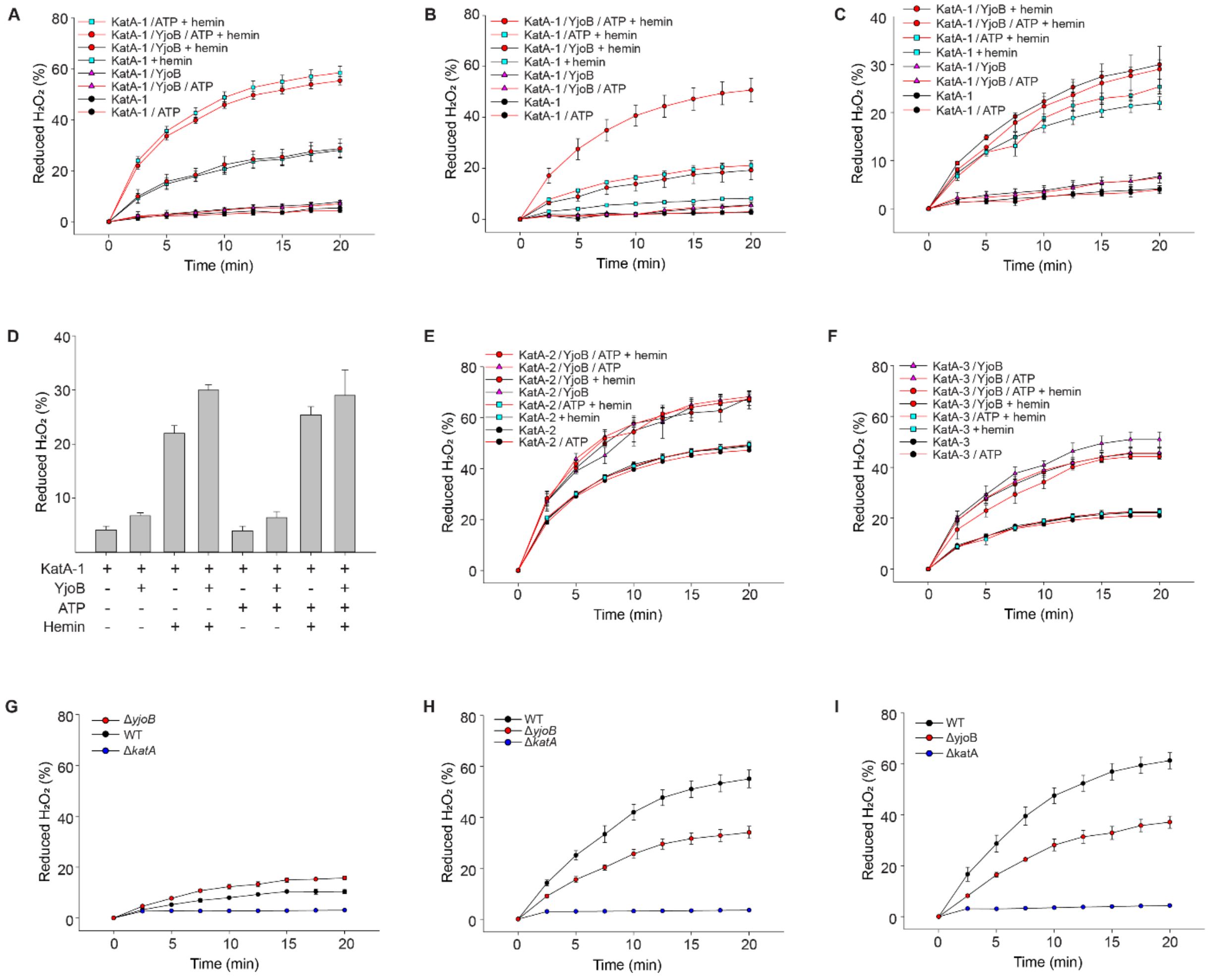

**Supplementary Figure S13. Catalase activity of KatA.** (**A, B**) Catalase activity of KatA-1. A mixture of KatA-1, YjoB, and ATP was incubated at 30 °C (A) and 42 °C (B) for 1 h and further incubated with or without hemin at 30 °C. The decomposition of $H_2O_2$ by KatA-1 was measured at 30 ˚C. The reaction mixture contained 25 nM KatA-1, 25 nM YjoB, 25 nM hemin, 5 μM ATP, and 20 mM $H_2O_2$. (**C**) Catalase activity of renatured KatA-1. KatA-1 denatured with 6 M guanidine hydrochloride was renatured by diluting in buffer A containing YjoB and ATP. $H_2O_2$ decomposition was measured after incubating with or without hemin for 30 min at 30 °C. The reaction mixture contained 25 nM KatA-1, 25 nM YjoB, 25 nM hemin, 5 μM ATP, and 20 mM $H_2O_2$. (**D**) Bar diagrams showing the level of $H_2O_2$ decomposition at the 20 min time point of (c). (**E, F**) Catalase activity of KatA-2 (E) and KatA-3 (F). The mixture of KatA-2 (or KatA-3), YjoB, and ATP was incubated at 42 ˚C for 1 h and further incubated with or without hemin at 30 °C. $H_2O_2$ decomposition was measured after incubating with or without hemin for 30 min at 30 °C. The reaction mixture comprised 0.25 nM KatA-2, 0.25 nM KatA-3, 25 nM YjoB, 25 nM hemin, 5 μM ATP, and 20 mM $H_2O_2$. (**G–I**) Decomposition of 20 mM $H_2O_2$ by cell lysates prepared in the exponential (G), early stationary (H), and extended stationary (I) growth phases of wild-type *B. subtilis* and its mutants (Δ*katA* and Δ*yjoB*). Wild-type *B. subtilis* and its mutants (Δ*katA* and Δ*yjoB*)

were collected at exponential phase ($OD_{600}$ = 0.65), stationary phase ($OD_{600}$ = 1.0), and 2 h after reaching stationary phase. Data are presented as the mean ± standard error of three replicates.

**Supplementary Figure S14**

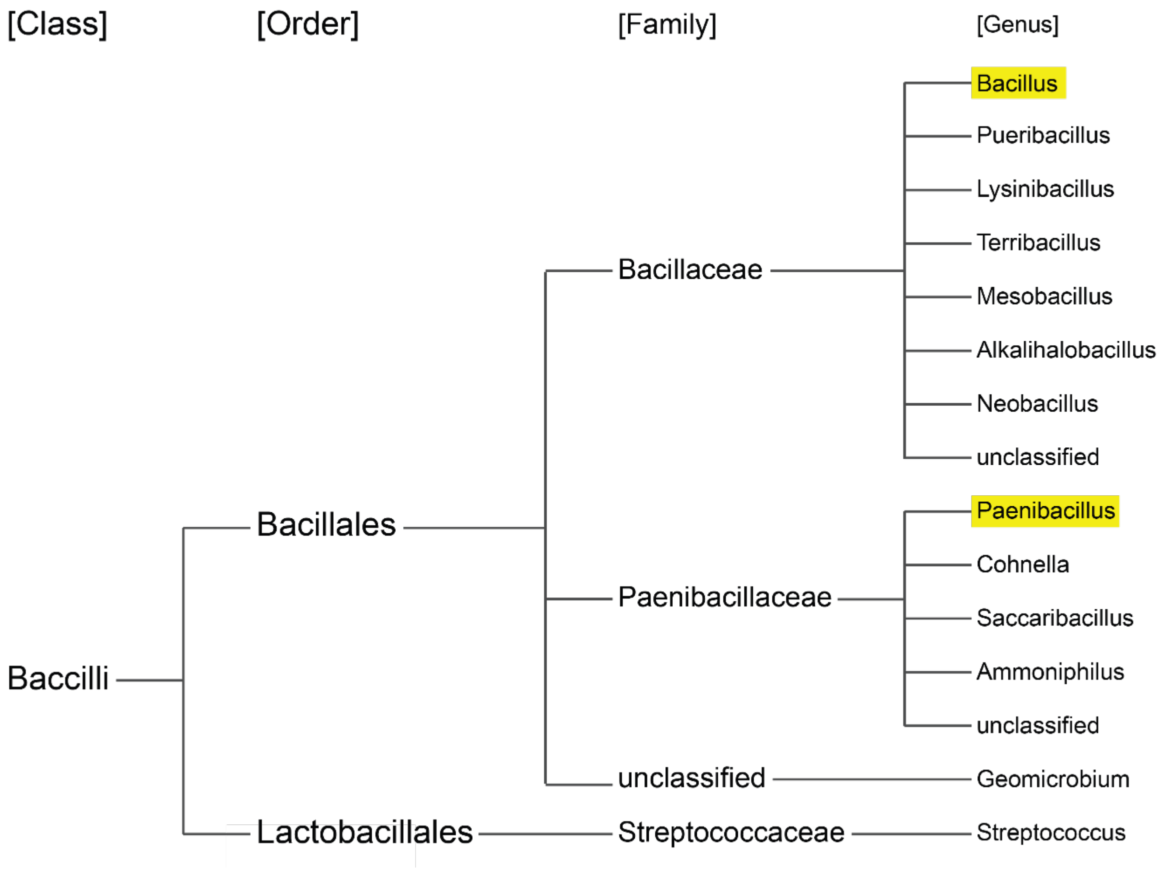

**Supplementary Figure S14. Phylogenic tree of bacteria harboring YjoB homologs.** The homologs of *B. subtilis* YjoB with more than 50% sequence identity were searched in the non-redundant protein sequences using BLAST. The phylogenic tree shows the genus with the YjoB homolog. The YjoB homologs are commonly found in the genera *Bacillus* and *Paenibacillus* (yellow boxes).

### Supplementary Figure S15

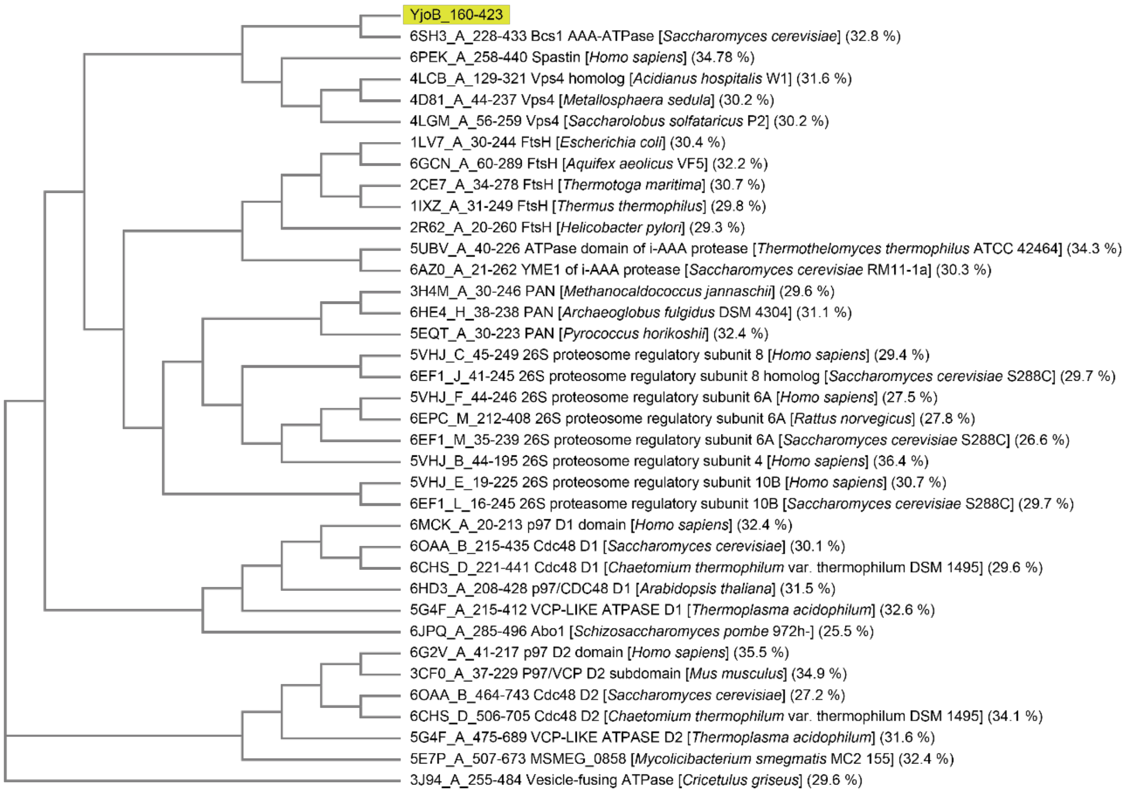

**Supplementary Figure S15. Phylogenic tree of AAA domains with known structures.** Homologs of YjoB-AAA (residues 160–423) were searched in the Protein Data Bank using BLAST. The phylogenic tree was prepared after multiple sequence alignment of the homologs.

### Supplementary Figure S16

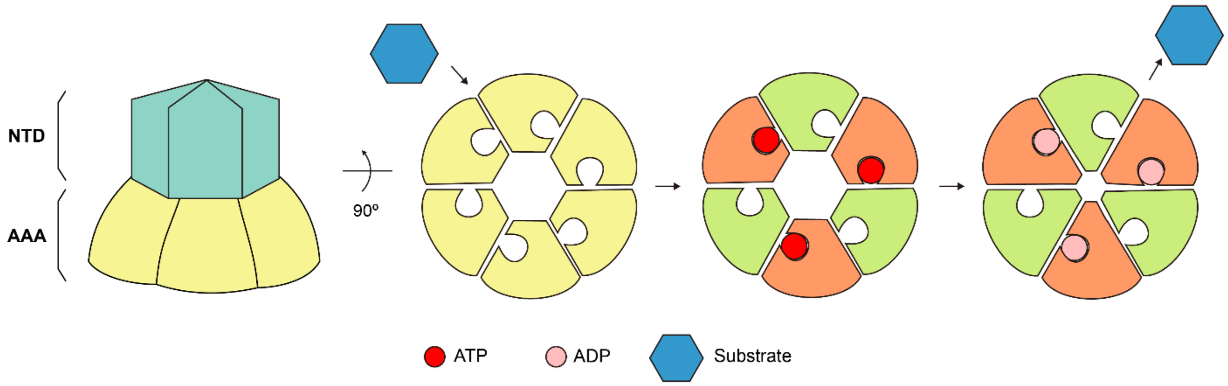

**Supplementary Figure S16. A model of nucleotide-dependent substrate binding.**

**Supplementary Table S1. Data-collection and refinement statistics for the structure determination.**

| **Data collection** | | | | | |
|---|---|---|---|---|---|
| **Data set** | YjoB | YjoB-ADP | MAD data Peak | MAD data Remote | YjoB-NTD |
| X-ray source | PLS II-BL11C | | | | |
| Space group | R32 | | | | C2 |
| Unit cell | | | | | |
| a, b, c (Å) | 155.6, 155.6, 303.1 | 156.3, 156,3, 302.8 | 155.1, 115.1, 302.9 | 155.3, 155.3, 303.2 | 144.3, 84.2, 180.2 |
| α, β, γ (°) | 90, 90, 120 | 90, 90, 120 | 90, 90, 120 | 90, 90,120 | 90, 112.3, 90 |
| Resolution (Å) | 50.0 – 2.62 (2.72 - 2.62) | 50.0 – 2.70 (2.81 – 2.70) | 50.0 – 2.90 (3.06 – 2.90) | | 50.0 – 3.00 (3.05 – 3.00) |
| Wavelength (Å) | 0.9793 | 0.9793 | 0.9794 | 0.9716 | 0.9794 |
| Total/Unique reflections | 464,860/42,649 (43,416/4,419) | 435,100/39,371 (48,981//4,361) | 343,774/31,382 (47,273/4,522) | 346,312/31,502 (49,296/4,566) | 124,952/39,002 (6,223/1,935) |
| Completeness (%) | 99.8 (99.5) | 100.0 (100.0) | 99.9 (100.0) | 100.0 (100.0) | 96.9 (97.7) |
| I/σ | 16.6 (5.8) | 16.2 (3.8) | 16.4 (7.8) | 19.5 (8.2) | 11.4 (1.5) |
| $R_{merge}$ (%) | 9.7 (34.2) | 12.0 (68.9) | 10.3 (24.7) | 9.3 (26.3) | 12.1 (70.8) |
| $R_{pim}$ (%) | 3.1 (11.4) | 3.8 (21.7) | 3.3 (7.9) | 2.9 (8.3) | 7.9 (45.7) |
| Figure of merit | | | 0.51 | | |
| **Refinement** | | | | | |
| Resolution | 50.0 -2.62 | 50.0 – 2.70 | | | 49.36 – 3.00 |
| No. reflections, working/free | 42,645/2,244 | 39,295/1,940 | | | 36,701/1,984 |
| $R_{work}/R_{free}$ (%) | 20.0/23.6 | 18.7/22.3 | | | 23.7/28.0 |
| No. atoms | 6,734 | 6,669 | | | 15,526 |
| RMSD | | | | | |
| Bond length (Å) | 0.008 | 0.009 | | | 0.003 |
| Bond angle (°) | 0.981 | 0.995 | | | 0.644 |
| B factors | 46.5 | 48.5 | | | 40.8 |
| Ramachandran plot (%) | | | | | |
| Favor | 96.7 | 96.0 | | | 96.5 |
| Allowed | 3.0 | 3.9 | | | 3.4 |
| Disallowed | 0.1 | 0.1 | | | 0.1 |
| Clashscore | 9.11 | 7.30 | | | 6.62 |

[a] Values in parentheses are for the highest resolution shell.

**Supplementary Table S2. Interactions between subunits in YjoB hexamer**

| Chain A | Chain B | Chain D | Chain E | Chain C |
|---|---|---|---|---|
| Pro5 | | | (H)Tyr76 | |
| Phe6 | | | (H)Arg31, (ф)Tyr76, (ф)Glu79 | |
| Ile7 | | | (ф)Leu28, (H)Arg31, (ф)Arg31, (ф)Thr35, (ф)Ala75, (H)Tyr76, (ф)Ile77 | |
| Tyr8 | (H)Ala19, (ф)Tyr22 | | (ф)Tyr27, (H)Ile77, (ф)Phe78 | |
| Gln9 | | | (H)Tyr27, (ф)Arg31 | |
| Tyr10 | (ф)Asn15, (ф)Arg17, (ф)Ala18 | | | |
| Glu12 | (ф)Gly23 | | | |
| Asn15 | | (H)Tyr10, (H)Lys13 | | |
| Arg17 | (H)Thr24, (ф)Tyr27 | | | |
| Ala18 | (H)Thr24 | (ф)Tyr10 | | |
| Ala19 | (ф)Phe78 | (H)Tyr8 | | |
| Tyr22 | (ф)Phe78, (H)Lys80 | (ф)Tyr8 | | |
| Thr24 | | (H)Arg17, (H)Ala18 | | |
| Phe25 | (ф)Leu81, (ф)Phe123 | | | |
| Gly26 | (ф)Lys80 | | | |
| Tyr27 | | (ф)Arg17 | | (ф)Tyr8, (H)Gln9 |
| Leu28 | | | | (ф)Ile7 |
| Arg31 | | | | (H)Phe6, (H)Ile7, (ф)Ile7, (ф)Gln9 |
| Glu33 | (H)Lys80 | | | (ф)Ile7 |
| Thr35 | | | | |
| Tyr47 | (ф)Lys84, (ф)Val99 | | | |
| Ser49 | (H)Glu82, (H)Arg104, (ф)Ile128 | | | |
| Asn53 | (H)Tyr88 | | | |
| Thr54 | (H)Glu90 | | | |
| Glu55 | (H)Tyr88, (ф)Glu90, (H)Glu90 | | | |
| Tyr56 | (ф)Tyr88 | | | |
| Ala75 | | | | (ф)Ile7 |
| Tyr76 | | | | (H)Pro5, (ф)Phe6, (H)Ile7 |
| Ile77 | | | | (ф)Ile7, (H)Tyr8 |
| Phe78 | | (ф)Ala19, (ф)Tyr22 | | (ф)Tyr8 |
| Lys80 | | (H)Tyr22, (ф)Gly26, (H)Glu33 | | |
| Leu81 | | (ф)Phe25, (ф)Ser49 | | |

| | | |
|---|---|---|
| Glu82 | | (H)Ser49 |
| Lys84 | | (H)Glu45, (ф)Tyr47, (H)Asp139 |
| Phe86 | | (ф)Leu134, (ф)Met142, (ф)Leu146 |
| Tyr88 | | (H)Asn53, (ф)Leu146 |
| Glu90 | | (H)Asn53, (H)Thr54, (H)Glu55 |
| Asp91 | | (H)Gln267 |
| Pro95 | | (ф)Arg150 |
| Tyr97 | | (ф)Leu146, (ф)Glu147 |
| Val99 | | (ф)Tyr47, (ф)Asp139 |
| Arg104 | | (H)Ser49 |
| Tyr122 | (ф)Gly126 | |
| Phe123 | | (ф)Tyr22, (ф)Phe25, (ф)Phe131 |
| Gln124 | (ф)Asp125 | |
| Gly126 | | (ф)Tyr122 |
| Ile128 | | (ф)Ser49, (ф)Phe131 |
| Phe131 | (ф)Gly126, (ф)Ile128 | |
| Leu134 | (ф)Phe86 | |
| Asp139 | (H)Lys84, (ф)Val99 | |
| Met142 | (ф)Phe86 | |
| Leu146 | (ф)Phe86, (ф)Tyr97 | |
| Glu147 | (ф)Tyr97 | |
| Asp168 | | (H)Glu260 |
| Arg180 | | (H)Asp216 |
| Arg199 | (ф)Gln386, (H)Gln389 | |
| Gln203 | (ф)Leu385, (H)Gln389 | |
| Phe211 | (ф)His388 | |
| Tyr215 | (ф)Ile354, (H)Tyr380 | |
| Asp216 | (H)Arg180, (H)Arg350 | |
| Ile217 | (ф)Tyr380, (ф)Val381 | |
| Pro218 | (ф)Arg180 | |
| Arg221 | (H)Gue378 | |
| Glu255 | | (H)Ser259, (ф)Val289 |
| Phe256 | | (ф)Ser258 |
| Ser259 | (H)Glu255 | |
| Glu260 | (H)Asp168 | |
| Glu263 | (H)Lys93 | |
| Val289 | (ф)Glu255 | |

| | | |
|---|---|---|
| Phe292 | (ф)Thr254 | |
| Asn295 | (H)Glu280 | |
| Gly299 | (ф)Thr234 | |
| Ala300 | (ф)Thr234,<br>(ф)Tyr250, (H)Glu280 | |
| Thr301 | (ф)Val163 | |
| Lys303 | (H)Glu176 | |
| Glu304 | (H)Arg180 | |
| Arg350 | | (H)Asp216 |
| Ile354 | | (ф)Tyr215 |
| Glu378 | | (ф)Arg325, (H)Ala326 |
| Tyr380 | | (H)Tyr215 |
| Ala384 | | (ф)Phe211 |
| Leu385 | | (ф)Phe211 |
| His388 | | (ф)Phe211, (ф)Tyr215 |
